# Two dominant axes structure high-dimensional object representations in the human ventral temporal cortex

**DOI:** 10.64898/2026.09.09.749976

**Authors:** Yun Wang, Kaixiang Zhuang, Xinyu Liang, Jianfeng Feng, Martin Hebart, Deniz Vatansever

## Abstract

The ventral temporal cortex is critical for visual object recognition, yet the principles organising its representations remain unclear. Using ultra-high-field 7T fMRI and dense sampling of 1,854 object concepts, we applied exploratory factor analysis to multivoxel response patterns to estimate the main axes of representational variation. Despite high dimensionality, over half of the explainable variance concentrated along two dominant axes: a continuum spanning objects of high biological salience, and a second integrating spatial context, physical scale, and manipulability. Importantly, neither reduced to a single semantic label, instead reflecting mixtures of visual and conceptual properties. Together they defined a triangular geometry, anchored by animal-, furniture- and food-related objects, stable across participants and two independent datasets, that mapped onto a lateral-to-medial cortical gradient, with category-selective regions emerging as local peaks within a broader continuum. These findings reveal that high-dimensional object representations in the ventral temporal cortex are structured by a small number of interpretable axes.

## Introduction

Understanding how the human visual cortex organises the wide range of objects encountered in everyday life constitutes a fundamental challenge in cognitive and computational neuroscience. Converging evidence from decades of research has established that the ventral temporal cortex plays a critical role in transforming visual inputs into representations that support object recognition and conceptual categorisation in both humans and non-human primates^1-3^. However, the principles that structure these representations remain contested.

A large body of research has long characterised this organisation primarily using pre-specified descriptors. Neuroimaging studies have mapped category-selective regions that respond preferentially to faces, bodies, scenes, words, and other ecologically salient object classes^4-6^, and lesion studies have revealed category-specific impairments following focal damage^7^. In parallel, multivoxel pattern decoding and representational similarity analyses have shown that information about object identity and category is distributed broadly across the ventral temporal cortex, extending beyond the boundaries of any single region^8-10^. Building on these findings, a growing body of research now describes object representations in the ventral temporal cortex in terms of continuous feature dimensions. For example, neural responses have been shown to vary smoothly along features such as animacy and real-world size^11-13^, and in the primate inferotemporal cortex objects have been mapped onto a continuous object space defined by a small number of pre-Wang et al. 2026 (preprint) 1 specified axes^14^. Such findings suggest that category-selective areas may reflect local peaks of sensitivity within a broader continuous representational space^15^, in which category boundaries emerge from graded variation across a multidimensional embedding.

Despite these recent advances, the precise nature of the dimensions that organise the wide array of object representations in the ventral temporal cortex remains unresolved. Most existing approaches test pre-defined models against neural data, linking ventral temporal cortex activity to semantic feature spaces^16^, behavioural semantic similarity judgements^17^, and deep neural network representations^18^. While highly informative, these approaches inherit assumptions embedded in their respective model spaces. As such, it remains unclear whether they reflect organisational principles intrinsic to the ventral temporal cortex population code, or whether alternative latent structures might provide a more accurate account of the underlying neural organisation.

A complementary and largely unexplored strategy is to infer latent dimensions directly from neural population responses. In this framework, ventral temporal cortex responses can be conceptualised as forming a high-dimensional representational manifold, with each object occupying a point defined by its multivoxel activity pattern^9,19^. Since neural population codes in this brain region are typically high-dimensional^20^, identifying the dominant axes along which object representations vary within this space would provide a principled characterisation of its large-scale organisation, independent of external semantic or behavioural labels^21^. This data-driven approach aligns with modern representation learning frameworks, in which latent variables are inferred from observations and interpreted post hoc without a priori definition^22^. However, two main practical obstacles have thus far constrained the use of such approaches to examine the neural architecture underlying object representation in the human ventral temporal cortex. First, existing neuroimaging studies have typically relied on relatively small stimulus sets that sparsely sample the multi-dimensional object space, limiting robust estimation of the representational geometry. Second, conventional fMRI affords limited spatiotemporal resolution, constraining how precisely population-level structure can be characterised. Overcoming both requires dense sampling across a broad range of object concepts together with high-fidelity measurements.

Here, we addressed these challenges by combining ultra-high-field 7T fMRI with dense sampling of 1,854 object concepts drawn from the THINGS image database^23^. Participants completed a continuous object recognition task across multiple scanning sessions, generating high-resolution neural responses to a broad sample of the object space. We then applied exploratory factor analysis to multivoxel neural response patterns to infer the dominant axes organising object representations in the ventral temporal cortex, without a priori model specification. Our findings revealed that, although this neural code was high-dimensional, its large-scale organisation was dominated by a small number of robust, interpretable axes that captured both classical categorical distinctions and graded variation across the object space. While these axes related to properties such as animacy and physical scale that have been proposed to organise object representations, they were broader than, and not fully explained by, any single pre-specified category or dimension. The axes generalised across participants, stimulus sets, and independent datasets, and mapped onto a lateral-to-medial cortical gradient spanning the ventral temporal cortex. Together, these results characterise the ventral visual stream as a high-dimensional representational code whose dominant axes are interpretable and recoverable directly from neural data, with relevance to both biological vision and computational theories of representation learning.

## Results

In order to uncover the latent organisation of object representations in the ventral temporal cortex using a data-driven approach, we combined ultra-high-field 7T fMRI with dense sampling of the THINGS image database (Fig. 1a). The stimulus set comprised 22,256 high-quality naturalistic object images spanning 1,854 concepts, upon which category-level analyses were based using 960 WordNet-labelled concepts grouped into 14 high-level semantic categories (Table S1)^23^. In total, twenty participants completed a continuous recognition task (old/new judgement) across five 7T scanning sessions, generating neural responses to 1,104 unique and 176 shared images per participant (Fig. 1b, c). Each image was presented three times in a pre-defined order^24^, and trial-wise response amplitudes were estimated using a robust analysis pipeline^25^. A ventral temporal cortex region of interest (ROI) was defined using a comprehensive framework incorporating functional localiser data, literature-based masks, and atlas-based parcellation schemes (Fig. 1d).

**Figure 1.**
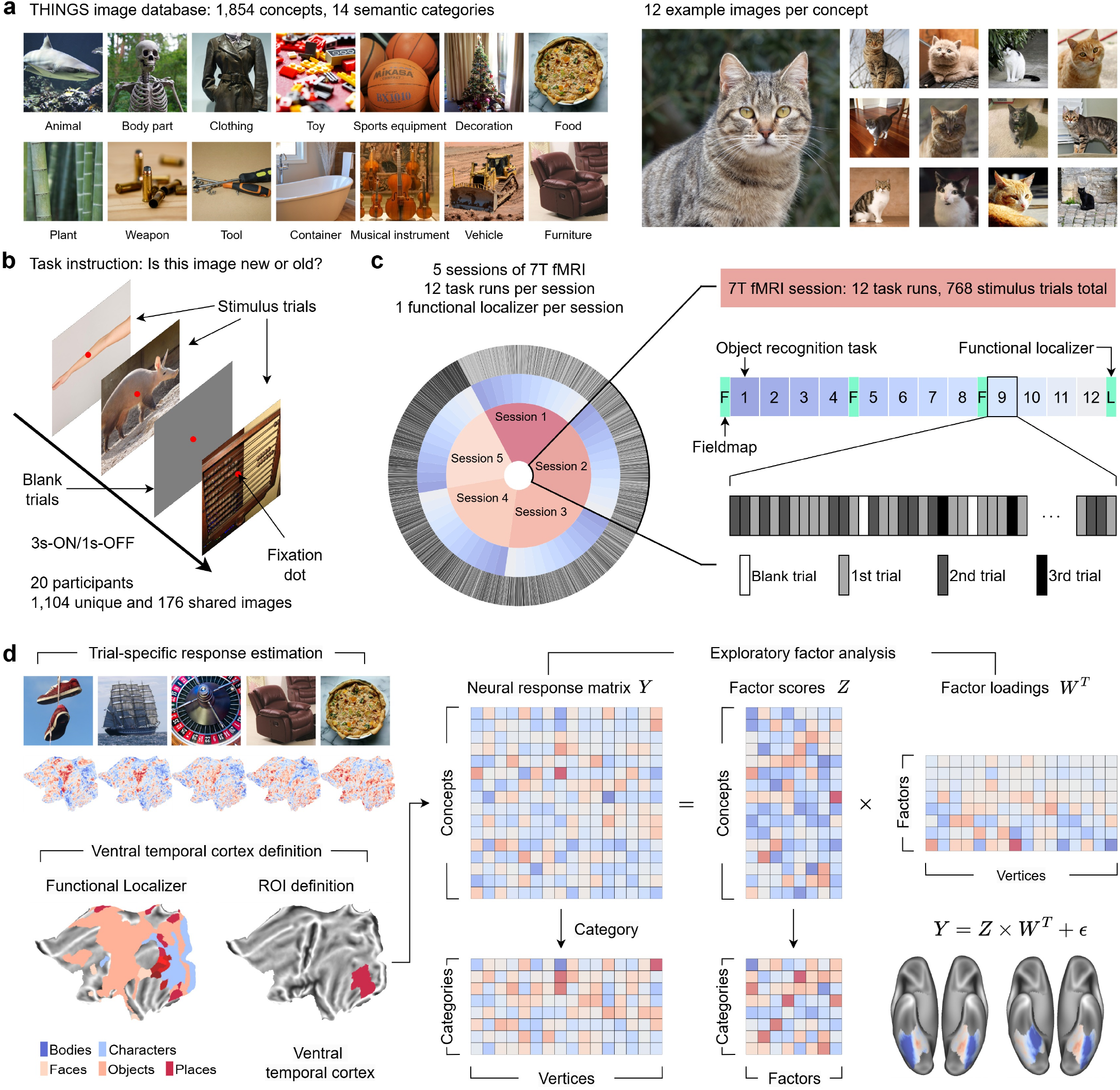
Experimental design and analytical framework. **a**. Experimental stimuli were drawn from the THINGS database, comprising 22,256 high-quality naturalistic object images spanning 1,854 object concepts, with category-level analyses based on 960 WordNet-labelled concepts organised into 14 high-level semantic categories. **b**. During each scanning session, 20 participants performed a continuous object recognition task, making old/new judgements. Participants viewed 1,104 unique and 176 shared images, each presented three times across the experiment. **c**. Every participant completed five 7T fMRI sessions on consecutive days. Sessions comprised 12 object recognition task runs (64 stimulus trials per run) and one functional localiser run. **d**. Trial-specific neural response amplitudes were derived using GLMsingle. The ventral temporal cortex region of interest (ROI) was defined as the union of functional localiser-based category-selective areas, literature-based ventral visual masks, and higher-order ventral visual parcels from the Human Connectome Project Multi-Modal Parcellation (HCP-MMP) atlas. Exploratory factor analysis was applied to the concept-by-vertex response matrix (Y), decomposing it into (i) latent factor scores (Z) capturing object-level representational structure and (ii) factor loadings (W) capturing vertex-level sensitivity profiles.

### Dense sampling of the object space elicits reliable neural responses

Our initial aim was to assess whether the employed dense sampling procedure generated reliable neural measurements of sufficient quality to estimate the representational geometry of the ventral temporal cortex. Behaviourally, participants’ task performance indicated that they remained actively engaged throughout the experiment (Fig. 2a). Response rates were high (mean = 97.81%, SD = 3.44%), and did not show a significant change across sessions (LMM likelihood-ratio test; *X*^2^(4) = 4.01, p = 0.40). In addition, adjusted hit rates (hit rate minus false alarm rate; mean = 0.47, SD = 0.23) remained above chance for the full duration of the experiment (one-sample t-test: t(19) = 10.38, p < 0.001, Cohen’s d = 2.32), confirming sustained and accurate task performance.

**Figure 2.**
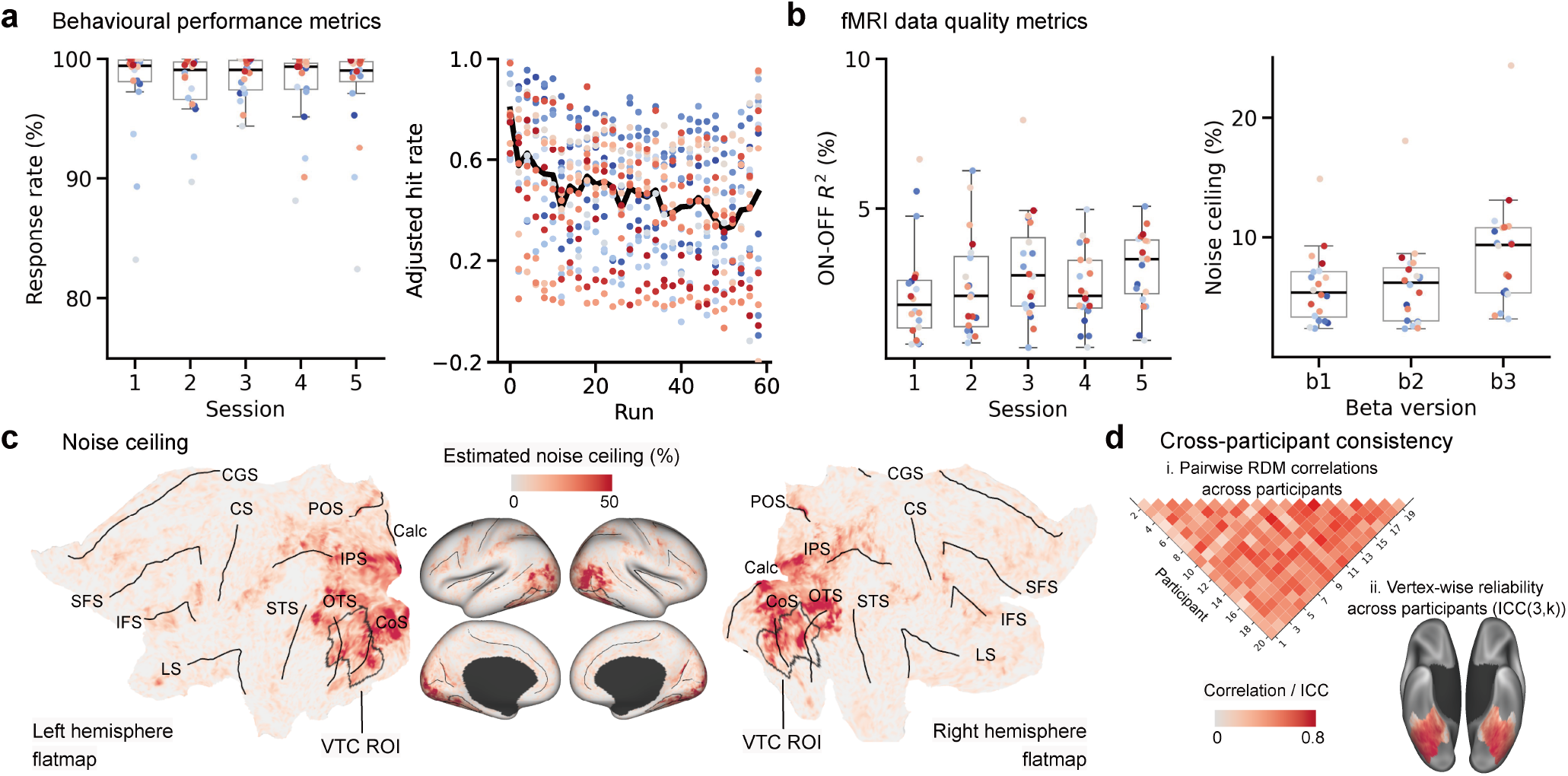
Dense sampling of the object space generates reliable behavioural and neural responses. **a.** Behavioural performance across sessions. Left: response rates per session, with each dot representing one participant. Right: adjusted hit rates (hit rate minus false alarm rate) across the experiment; the thick black line indicates the median across participants, and each dot represents a binned average across two consecutive runs. **b.** Task-evoked BOLD response reliability and noise ceiling within the ventral temporal cortex. Each data point denotes the median value per participant (n = 20). For all boxplots, the centre line denotes the median, box bounds the 25th and 75th percentiles, and whiskers extend to 1.5 times the interquartile range. Beta estimates are labelled as follows: b1, canonical haemodynamic response function (HRF); b2, voxel-wise HRF fitted using a library-of-HRFs approach; b3, library-of-HRFs approach with additional GLMdenoise and ridge regression. **c.** Noise ceiling map for a representative participant, with the ventral temporal cortex outlined in black. Anatomical landmarks: Calc, calcarine sulcus; CGS, cingulate sulcus; CoS, collateral sulcus; CS, central sulcus; IFS, inferior frontal sulcus; IPS, intraparietal sulcus; LS, lateral sulcus; OTS, occipitotemporal sulcus; POS, parieto-occipital sulcus; SFS, superior frontal sulcus; STS, superior temporal sulcus. **d.** Cross-participant consistency in neural responses to the 176 shared images. Left: pairwise Pearson correlations between category-level representational dissimilarity matrices (RDMs) across all participant pairs. Right: vertex-wise response reliability across participants, quantified using the intraclass correlation coefficient ICC(3,k) with k = 20.

Consistent with these behavioural observations, the 7T task fMRI data were also of high quality. Temporal signal-to-noise ratio was consistently strong (tSNR; overall mean = 55.32, SD = 16.61), and head motion was low (overall mean framewise displacement = 0.12 mm, SD = 0.12) across participants and sessions (Fig. S1). Moreover, average noise ceiling in the ventral temporal cortex was consistent across participants (overall mean = 8.72%, SD = 4.74%), and ON-OFF R^2^ values were elevated and stable across sessions (overall mean = 2.59%, SD = 1.55%; Fig. 2b-c), together indicating robust, reproducible task-evoked responses.

With the aim of quantifying cross-participant consistency, we focused on the 176 images shared across all participants. Vertex-wise responses to these images were highly reliable, with a median intraclass correlation coefficient of ICC = 0.63 (mean = 0.56, SD = 0.23; Fig. 2d; Fig. S2). Pair-wise correlations between high-level category representational dissimilarity matrices (RDMs) showed reproducible representational structure across individuals (back-transformed Fisher-z mean Pearson correlation r = 0.41, 95% participant-cluster bootstrap CI [0.35, 0.46]). Within-participant split-half analyses resulted in a mean Spearman-Brown-corrected RDM reliability of R = 0.34 (SD = 0.16). Standard RSA lower and upper noise ceilings were 0.62 and 0.67, respectively. Together, these results establish that the dense sampling approach produced a stable, high-fidelity neural dataset suitable for characterising the dominant organising principles underlying visual object representation in the human ventral temporal cortex.

### Two dominant axes capture most of the variance in object representations

With data quality established, we next examined the organisation of object representations in the ventral temporal cortex. For improved interpretability, we focused our main analyses on the 960 WordNet-labelled concepts with non-overlapping image sets. Trial-wise estimates were averaged to produce image-level responses, which were then aggregated within concepts to derive concept-level neural response patterns. In order to characterise the geometry of these responses, we applied exploratory factor analysis, which has previously been shown to successfully identify interpretable latent dimensions that capture shared variance across high-dimensional data^26,27^. Moreover, the concept response matrix was well suited to factor analysis, as indicated by a Kaiser-Meyer-Olkin measure of 0.96 and a significant Bartlett’s test of sphericity (p < 0.001), confirming that the data contained substantial shared covariance amenable to factor decomposition.

Split-half cross-validation showed that held-out log-likelihood peaked at 50 factors (Fig. S3a), indicating that the ventral temporal cortex code is high-dimensional and carries fine-grained representational detail, consistent with the high dimensionality reported for neural population responses more generally^20,21^. Within this solution, however, the first two factors were strongly dominant, together accounting for more than 50% of the variance and showing substantially greater stability than higher factors (Figs. S3b-c). In line with this observation, the recomputed category-level RDM derived from the two-dimensional factor scores correlated substantially with that derived from the full vertex-wise data (Pearson’s r = 0.60, 95% category-node bootstrap CI [0.31, 0.84]; Spearman-Brown-corrected neural-RDM reliability R = 0.93; conditional fixed-reference ceiling = 0.96; 62.5% of the ceiling), indicating that the two axes preserved core representational structure of the ventral temporal cortex. We therefore focused our subsequent analyses on this two-dimensional solution, while noting that later factors carry reliable, finer-grained information^28^.

When projected into this two-dimensional factor space, object representations formed a distinct triangular structure (Fig. 3a), in which the 14 high-level semantic categories occupied systematically different positions. Animal-related, food-related, and furniture-related objects anchored the three poles of the triangle, while other categories fell along the edges and interior, producing a continuous and overlapping representational map. Consistent with this smooth organisation, the Silhouette score was slightly negative (-0.12), indicating that categories blend into one another and do not reflect sharp, discrete clusters. Nevertheless, despite this graded organisation, the three-way separation among objects within animal, food, and furniture categories at the three distal poles^29-33^ was sufficiently pronounced to support high classification accuracy (86.9%, linear Support Vector Machine, Welch’s t-test against a label-shuffled baseline, p < 0.001).

**Figure 3.**
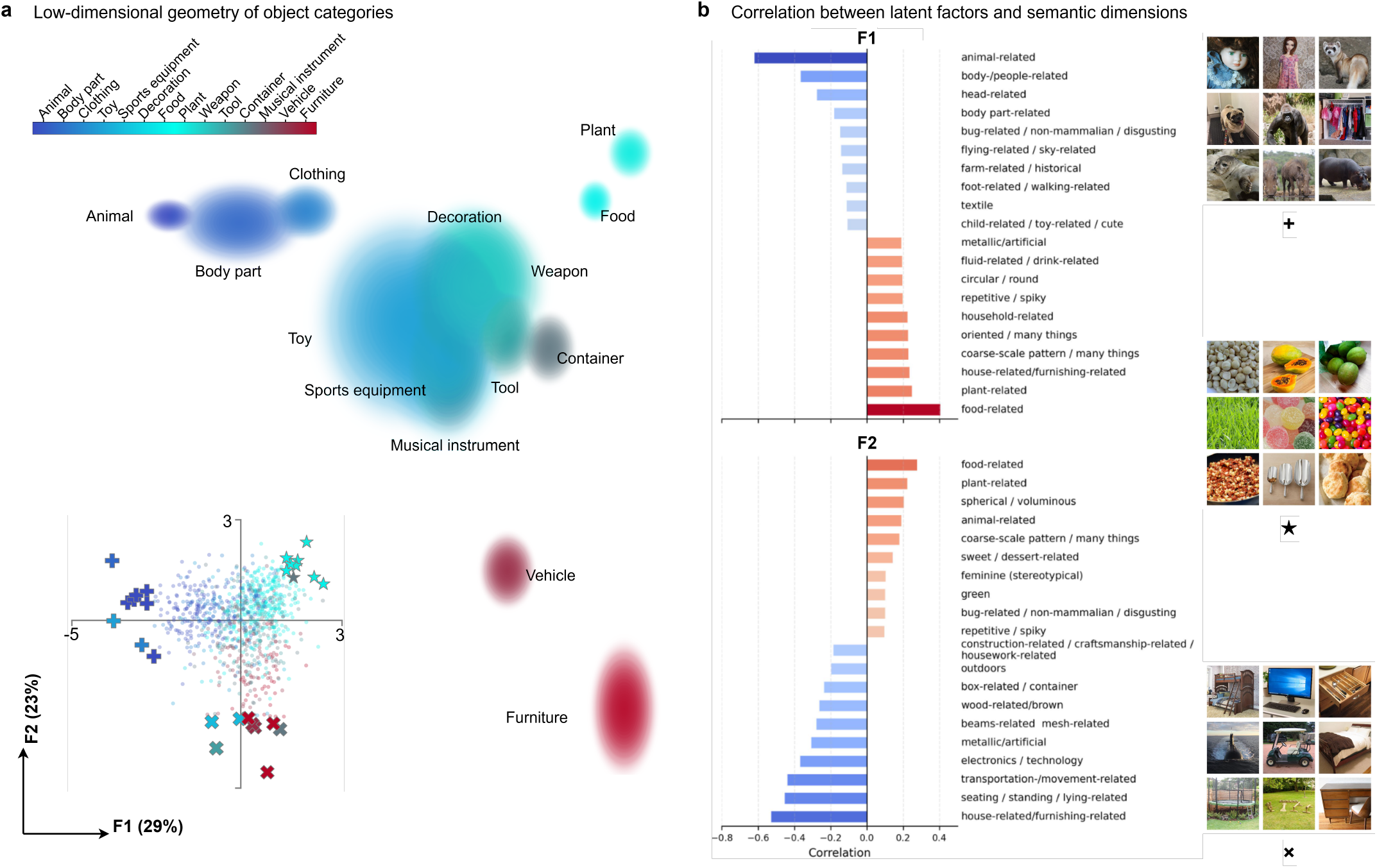
Object representations in the ventral temporal cortex are structured by a dominant and continuous two-dimensional geometry. **a.** Projection of object representations onto the two-dimensional factor space defined by Factor 1 and Factor 2. Upper panel: category-level distributions for 14 high-level semantic categories, with shaded regions indicating 95% confidence intervals. Lower panel: individual object concepts (n = 960) coloured by superordinate category; concepts at the three poles of the triangular geometry are marked by +, ×, and ★, with representative images shown on the right panel. **b**. Behavioural grounding of the latent factors. Bar plots show Pearson correlation coefficients between each factor score and 66 continuous semantic dimensions derived from independent human similarity judgements^42^. For each factor, the 10 dimensions with the largest positive and largest negative correlations are displayed, ranked by correlation magnitude. For privacy protection, the original photograph representing the face concept (top-left image in the exemplar grid) was replaced with a doll-face exemplar from the THINGS image database. The substitution is limited to figure visualization only.

The two axes of this geometry captured distinct and ecologically meaningful organisational principles. Factor 1 broadly tracked an animacy-related continuum with high biological salience, but also drew in inanimate objects with distinctly human-like features such as dolls, masks, and wigs (Fig. S4a). This mirrors prior evidence that face-like and schematic stimuli engage the same neural pathways as real faces and biological agents^34-36^ and is consistent with animacy being represented not as a single binary contrast but as a graded, multi-featured dimension^37,38^. Factor 2 separated furniture-related and vehicle-related objects from the rest of the object space. These categories shared strong associations with indoor and outdoor scenes, but also co-varied with real-world object size and manipulability^39-41^. Large, non-manipulable objects clustered at one pole, while small, graspable objects occupied the other (Fig. S4b). As such, neither scene association nor size alone fully accounted for this axis, pointing to a much richer organisational dimension that integrates multiple object properties.

With the aim of behaviourally grounding these interpretations in independent evidence, we correlated the factor scores with 66 continuous dimensions derived from large-scale human similarity judgements^42^, which provide a data-driven, interpretable description of the object properties that structure human similarity behaviour. Factor 1 showed its strongest negative correlations with animal-related, body-/people-related, and head-related dimensions, consistent with the placement of animal- and body-related objects at one pole. Factor 2 showed its strongest negative correlations with house-related/furnishing-related, seating-related, and transportation-/movement-related dimensions, consistent with the placement of furniture and vehicles at the negative pole of this axis (Fig. 3b, Fig. S5). Interestingly, both factors shared a common positive correlation associated with food-related and plant-related object dimensions, dominated by fruits and vegetables (Fig. S4c). Together, these correspondences provide converging, behaviourally-grounded support for the structure of this data-driven, neurally identified space.

In order to test whether established object dimensions could account for the two dominant axes, we used nested cross-validated ridge regression to predict Factor 1 and Factor 2 scores across the 960 concepts. While animacy and real-world size explained 38.7% and 29.8% of held-out variance, respectively, the 66 continuous dimensions explained 61.4% and 48.5% with the combination of the two predictor sets achieving 61.6% and 50.0% explained variance. We then tested whether the factor-derived representational geometry improved participant-held-out prediction of neural RDMs beyond the combined a priori models. Mean cross-validated R^2^ increased from 0.10 to 0.22 after adding the factor-derived RDM, corresponding to a mean ΔR^2^ of 0.12 that was positive in 96.0% of the train-test directions. Together, the results indicate that the recovered axes substantially overlapped with, but were not fully captured by the tested feature spaces, suggesting the presence of an organisational structure inherent to the neural data.

### Latent dimensions of object representation are stable and generalisable

We next asked whether the two-dimensional geometry we identified was robust to variation in stimulus sampling, experimental context, and dataset. For that purpose, we evaluated consistency at three distinct levels: category-level RDMs, the structure of the two-dimensional geometry, and factor loadings, while aligning factor order across comparisons using a factor-matching procedure.

First, we examined whether the geometry depended on the particular mixture of object categories in our stimulus set. Repeatedly constructing balanced datasets by sampling equal numbers of concepts from each category and refitting factor analysis produced solutions that were highly consistent with the full 960-concept results. Category-level neural RDMs from each category-balanced subset were strongly correlated with the full-data RDM (Fisher-z mean Pearson correlation r = 0.92, 95% bootstrap CI for the mean [0.91, 0.93]). After component matching and sign alignment, factor-loading correlations with the full-data solution were high for both dimensions (Factor 1: mean absolute Pearson’s r = 0.984, 95% bootstrap CI [0.983, 0.985]; Factor 2: r = 0.992, 95% bootstrap CI [0.9918, 0.9922]; Fig. 4a). The corresponding two-dimensional Euclidean-distance category RDMs were also highly similar to the full-data geometry (Fisher-z mean Pearson correlation r = 0.958, 95% bootstrap CI [0.955, 0.961]). These results indicate that the recovered geometry was robust to category-balanced resampling of concepts and was unlikely to be driven by unequal representation of any single category.

**Figure 4.**
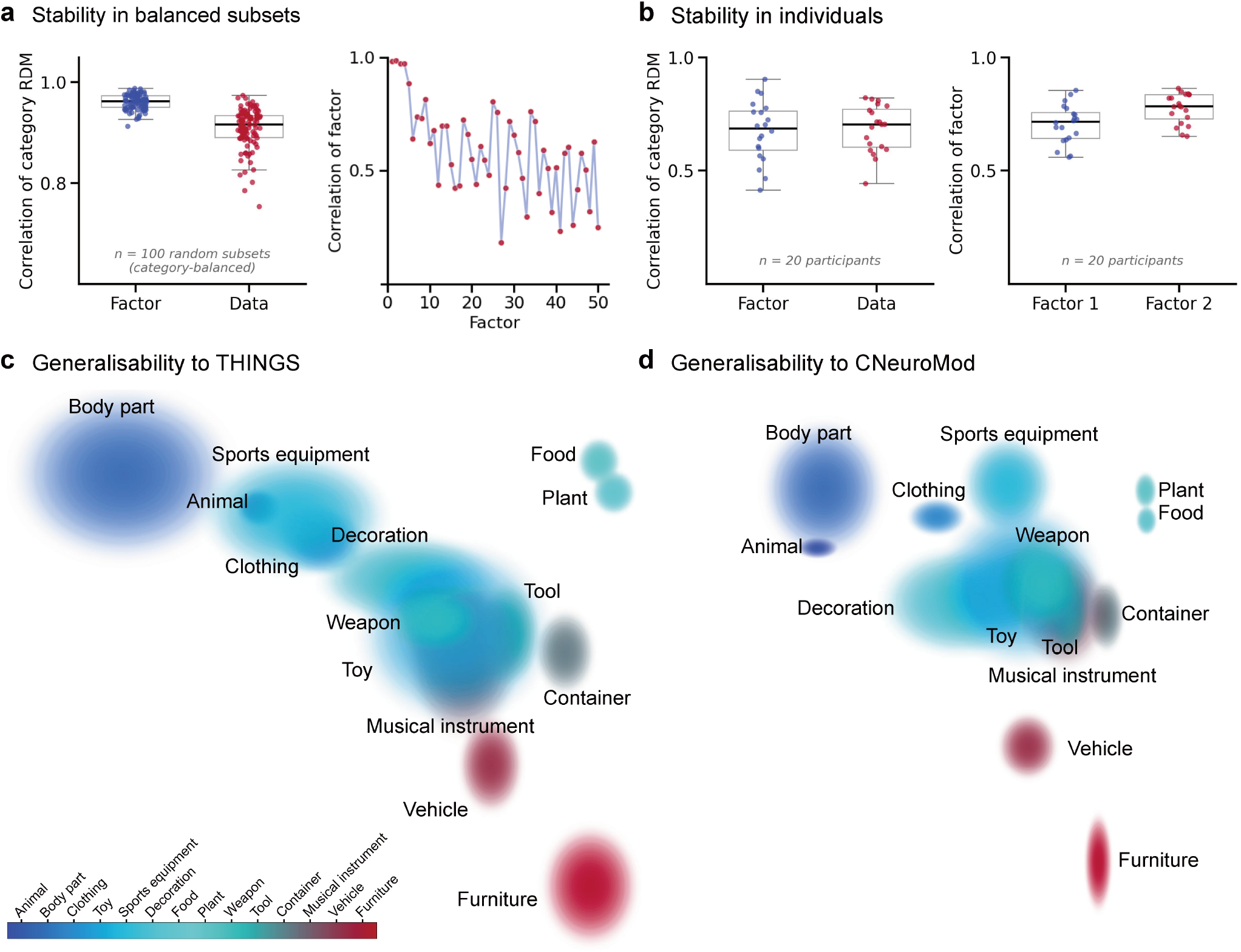
The two-dimensional representational geometry is robust and generalisable. **a.** Stability under balanced category sampling. Left: correlations between category-level RDMs from 100 category-balanced concept subsets (n = 20 concepts per category, 280 concepts total) and the full 960-concept dataset, computed in factor space (Euclidean distance; Factor) and vertex space (correlation distance; Data). Right: factor loading correlations between balanced subsets and the full dataset across the first 50 factors. **b.** Consistency across individual participants. Left: correlations between participant-level and group-level category RDMs in factor space and vertex space. Right: correlations between group-level and individual-participant factor loadings for Factor 1 and Factor 2 (n = 20 participants). **c-d.** Replication in independent datasets. The characteristic triangular geometry is reproduced by direct application of factor analysis to the THINGS-fMRI dataset (**c**) and the CNeuroMod-THINGS dataset (**d**), despite differences in participant samples, scanner field strength (3T versus 7T), and experimental design. Shaded regions indicate 95% confidence intervals for each superordinate category. The sign of Factor 2 is reversed relative to our dataset for visual alignment in panel d. Since factor sign is arbitrary, this does not alter the underlying representational geometry.

We then tested whether the latent space generalised to stimuli withheld from the main analysis. Responses to the 176 held-out shared images, projected into the factor space defined by the 960-concept solution, recovered a similar triangular structure (Fig. S6). The vertex-space correlation-distance category RDM correlated significantly with that from the full dataset (Pearson’s *r* = 0.59, 95% category-node bootstrap CI [0.14, 0.86]; permutation p < 0.001; conditional fixed-reference ceiling = 0.95; 62.3% of the ceiling). The two-dimensional Euclidean-distance category RDM obtained by projecting the held-out responses into the fixed factor space was similarly correlated with the full-data RDM (Pearson’s r = 0.73, 95% category-node bootstrap CI [0.49, 0.88]; permutation p < 0.001; ceiling = 0.95; 76.4% of the ceiling). Individual-participant solutions were also consistent. Factor analysis run separately on each of the 20 participants produced geometries closely matching the group-level result in factor loadings, vertex-space category RDMs, and two-dimensional factor-space Euclidean-distance category RDMs (Fig. 4b, Fig. S7).

Finally, we tested whether the same geometry emerged in entirely independent datasets. Applying our analysis pipeline to the THINGS-fMRI dataset^43^ and the CNeuroMod-THINGS dataset^44^, both acquired at 3T with different tasks and experimental designs, reproduced the characteristic triangular organisation (Figs. 4c-d). For THINGS-fMRI, the vertex-space correlation-distance category RDM correlated significantly with that from the full dataset at r = 0.84 (95% category-node bootstrap CI [0.56, 0.95]; permutation p < 0.001; conditional fixed-reference ceiling = 0.95; 88.4% of the ceiling). The matched two-dimensional factor-space Euclidean-distance category RDM correlation was r = 0.67 (95% category-node bootstrap CI [0.38,0.91]; permutation p < 0.001; conditional fixed-reference ceiling = 0.91; 73.2% of the ceiling), and the absolute factor-loading correlation across Factors 1 and 2 was r = 0.73 (Fig. S8). For CNeuroMod-THINGS, the corresponding vertex-space correlation-distance category RDM correlation was *r* = 0.80 (95% category-node bootstrap CI [0.56, 0.94]; permutation p < 0.001; conditional fixed-reference ceiling = 0.98; 81.3% of the ceiling), and the matched two-dimensional factor-space Euclidean-distance category RDM correlation was *r* = 0.91 (95% CI [0.76, 0.97]; permutation p < 0.001; conditional fixed-reference ceiling = 0.98; 92.8% of the ceiling). Absolute factor-loading correlations were r = 0.88 and r = 0.93 for Factors 1 and 2, respectively (Fig. S9). Taken together, across differences in scanner field strength, task design, and spatiotemporal resolution, the core category-level representational geometry was recovered in both independent datasets.

### Latent dimensions align with a lateral-to-medial cortical topography

Having established the structure and robustness of the two-dimensional geometry, we finally asked how this organisational architecture is implemented across the surface of the ventral temporal cortex. We projected factor loadings back onto the cortical surface and examined how high-level category responses embedded within the subspace varied across ventral temporal regions, thus testing whether the latent dimensions corresponded to a continuous topographic gradient.

The spatial distribution of factor loadings revealed a clear lateral-to-medial organisation (Fig. 5a). Factor 1 loadings were strongest in magnitude in lateral ventral temporal regions, overlapping substantially with the Fusiform Face Area (FFA), while weaker positive loadings extended more medially. Factor 2 showed the complementary pattern in that the strongest negative loadings fell in the medial ventral temporal cortex, overlapping with the Parahippocampal Place Area (PPA), while weaker positive loadings extended more laterally. Importantly, these distributions were not segregated into strictly discrete territories. Opposite-signed loadings and overlapping intermediate zones were present throughout, indicating that the two factors are embedded within a continuous, partially overlapping lateral-to-medial topography instead of two anatomically distinct patches.

**Figure 5.**
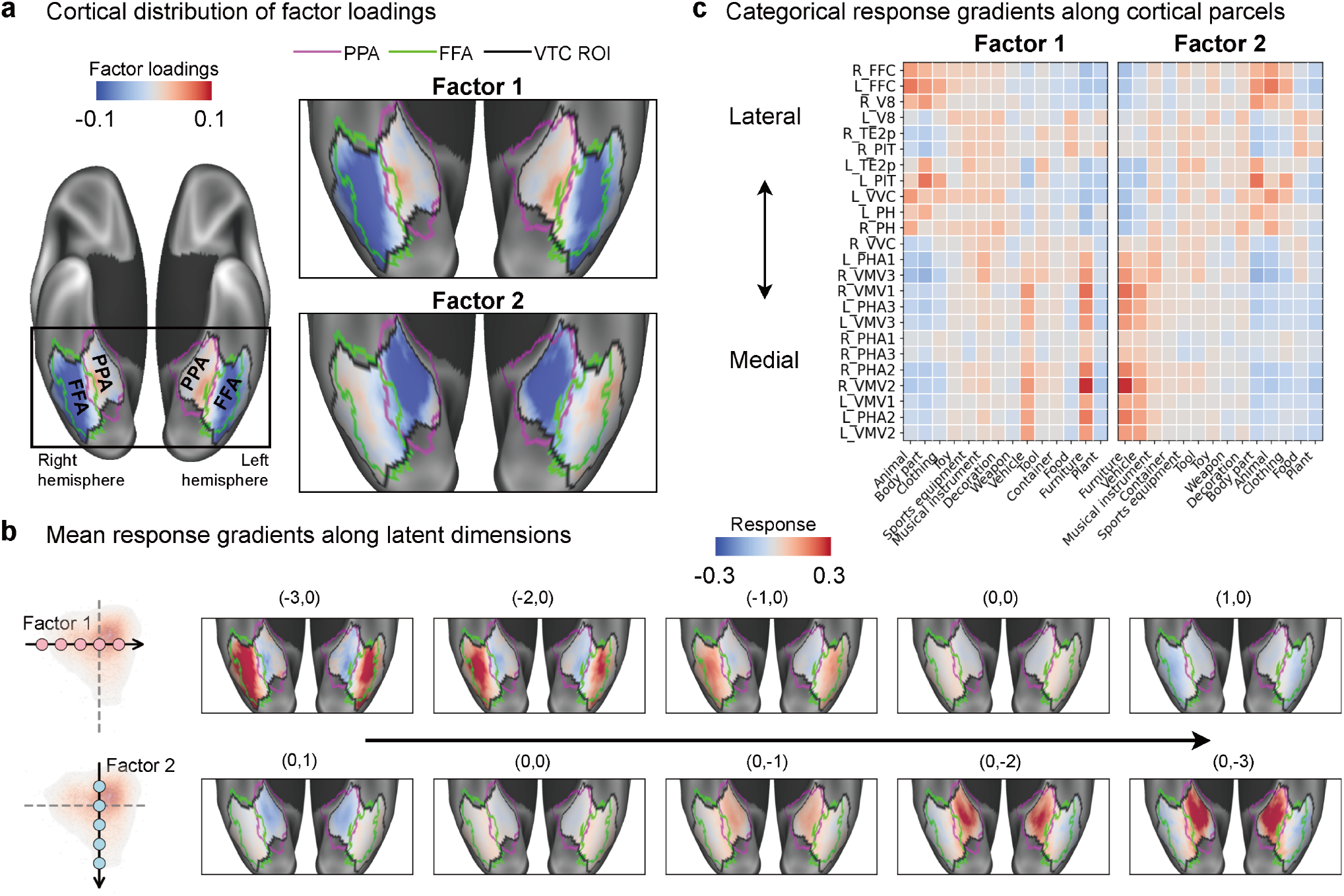
The two-dimensional latent geometry is neurally implemented as a smooth lateral-to-medial cortical gradient. **a.** Cortical distribution of factor loadings (W) for Factor 1 and Factor 2, projected onto ventral cortical surfaces. Factor 1 loadings are strongest in lateral ventral temporal regions; Factor 2 loadings are strongest in medial ventral temporal regions. Fuchsia and lime outlines denote the Fusiform Face Area (FFA) and Parahippocampal Place Area (PPA), respectively, defined from the Natural Scenes Dataset (NSD) functional localiser task^24^. **b**. Mean cortical response patterns sampled at selected coordinates along each latent dimension. Schematics indicate sampled positions within the two-dimensional factor space, with coordinates labelled above each surface map. Response profiles transition smoothly across both axes, shifting gradually between lateral- and medial-weighted patterns without abrupt boundaries. **c.** Category-level neural response gradients across HCP-MMP parcels within the ventral temporal cortex. Heatmaps show mean neural responses for 14 high-level categories across parcels ordered by a normalised FFA-PPA overlap index (proportion of parcel vertices overlapping the FFA localiser minus the proportion overlapping the PPA localiser). Left panel: categories ordered by mean Factor 1 score; right panel: categories ordered by mean Factor 2 score. Graded shifts in category responses across the parcel ordering reflect the continuous lateral-to-medial topographic organisation of the latent geometry.

This pattern was more evident when we sampled cortical response patterns at selected coordinates along each latent dimension (Fig. 5b). Along Factor 1, responses in lateral ventral temporal cortex were strongest at the most negative coordinates and declined progressively toward zero, giving way to a weak inverse profile at positive coordinates. Along Factor 2, medial responses around the PPA strengthened progressively from positive through zero to negative coordinates, peaking at the most negative point. Importantly, intermediate positions along both axes showed mixed lateral and medial contributions rather than abrupt transitions, confirming that traversing these dimensions is accompanied by smooth, gradual shifts in cortical response profile.

Moreover, category-level response gradients across ventral temporal parcels corroborated this observation (Fig. 5c). Ordering parcels by a normalised FFA-PPA overlap index, we found that responses to categories with large negative Factor 1 scores (e.g. animal-related, body part-related) were strongest toward the FFA-weighted end, while responses to categories with large negative Factor 2 scores (e.g. furniture-related, vehicle-related) were strongest toward the PPA-weighted end. Food- and plant-related objects, positioned between the poles of both dimensions, showed more distributed responses across the parcel ordering. A broadly similar gradient was evident in category decoding accuracy across parcels (Fig. S10).

Together, these results suggest that the two-dimensional latent geometry topographically maps onto a smooth lateral-to-medial cortical gradient spanning the ventral temporal cortex, while classical category-selective regions such as the FFA and PPA emerge as local response maxima within a broader topographic continuum.

## Discussion

A central challenge in visual and computational neuroscience is to identify the principles that organise object representations in the human ventral temporal cortex^1,2^. Combining ultra-high-field 7T fMRI with dense sampling of object concepts from the THINGS image database^23^, we show that the large scale organisation of the ventral temporal cortex code, although high-dimensional^20,21^, is dominated by two axes recoverable directly from neural population responses. These axes define a continuous triangular geometry, anchored by animacy-related and food-related concepts together with inanimate, scene-associated ones, that accounted for the main directions of variation, generalised across participants, stimulus sets and independent datasets, and topographically mapped onto a continuous lateral-to-medial cortical gradient. As this structure was derived directly from neural data, our findings establish that ecologically meaningful organisational principles are intrinsic to the human ventral temporal cortex population code.

The identified geometry converges with earlier accounts while also revising their central claims. Objects in the primate inferotemporal cortex have been mapped onto a continuous object space defined by a small number of pre-specified axes^14^, and the human ventral stream has been described as tripartite, dividing animate objects, large inanimate objects and small inanimate objects along animacy and real-world size^11,29,30^. Our geometry converges with both accounts in recovering a compact, three-poled description, and two of its poles align closely with the established scheme, in that animal- and body part-related concepts anchor one extreme of Factor 1 while furniture and vehicles anchor the negative extreme of Factor 2. The third pole, however, is not occupied by small manipulable objects. Tools, containers and other readily handled items instead fell along the edges and interior of the triangle, and the remaining extreme was occupied by food- and plant-related concepts. Selectivity for food in the human ventral visual cortex has only recently been described, initially through hypothesis-free decomposition of voxel responses^31^ and subsequently in independent samples using converging approaches^32,33^, and the extent to which these findings reflects a genuine organising principle has remained open. Here, food- and plant-related concepts anchored the geometry, and did so under category-balanced resampling, in held-out stimuli, in individual participants, and in two independent datasets. Since the plant category comprises largely fruits and vegetables, this pole may be better characterised in terms of edibility, and as such, food-related structure constitutes a reproducible property of the ventral temporal cortex code, situated within a more general organisational geometry.

Beyond the extreme poles, the two dominant axes each carried a clear and ecologically meaningful interpretation. The first factor aligned broadly with the animate-inanimate distinction, but its scope extended well beyond biological animacy to include inanimate objects with distinctly human-like features, such as dolls, masks, and wigs that were clustered alongside animals, consistent with evidence that face-like and schematic stimuli engage similar neural pathways to real faces^34,36^ and biological agents^35^. We therefore interpret Factor 1 as a graded continuum that also reflects perceived human-likeness or biological salience^37,38^, mapping naturally onto the established tuning of lateral ventral temporal cortex for faces, bodies, and socially relevant stimuli^5,45^, with the classical animate-inanimate distinction emerging as only one expression of this broader axis. The second dimension grouped furniture and vehicles at one pole, a pairing that initially appears to reflect shared associations with indoor and outdoor scenes^4,46^, yet it also co-varied with real-world object size^11,30^ and ease of manual manipulation^12,39^, separating large, non-manipulable objects from small, readily handled ones. The convergence of scene context, physical scale, and action affordance along a single axis suggests that Factor 2 integrates these properties into a unified representational dimension^40^.

This distribution of category-selective tuning within the identified two-dimensional geometry helps reconcile previously divergent observations. Categories such as animals and furniture occupy extreme positions along the two axes, producing regions of concentrated selectivity, whereas other categories fall along the edges and interior of the triangle in graded transitions. This arrangement may explain why univariate analyses reliably identify localised peaks of category selectivity^4,5,31,47^, while multivoxel pattern analyses decode object information across the full extent of the ventral temporal cortex^8,10,48,49^. It is possible that both of these observations follow from a single continuous space in which local categorical distinctions arise from graded variation along a few dominant axes^37,40^. Consistent with this view, our results suggest that the two-dimensional geometry was implemented as a smooth lateral-to-medial cortical gradient, with Factor 1 loadings strongest in lateral regions overlapping the FFA, and Factor 2 loadings strongest in medial regions overlapping the PPA. As such, traversing the manifold from pole to pole corresponded to moving across the cortical surface from lateral face-selective to medial scene-selective regions^4,5,45,46^, with response profiles shifting gradually across this axis. Classical category-selective regions thus emerge as local response maxima within a broader topographic continuum^15^, which represents a spatially efficient scheme in which fine-tuned selectivity arises at the poles of continuous representational gradients, and the localised and distributed descriptions of ventral temporal cortex organisation become complementary readouts of the same underlying structure.

That two axes dominate a code that is otherwise high-dimensional does not by itself follow from variance concentration. Cross-validated model selection favoured a 50-factor solution, and population responses in the visual cortex occupy many reliable dimensions with approximately scale-free eigenspectral^20,21,28^, under which some concentration of variance in the leading directions is expected on statistical grounds alone. The informative observation is therefore not merely the proportion of variance captured, but that the two dominant axes proved interpretable against independent behavioural dimensions^42^, stable across resampling, individuals and datasets, and organised as a smooth cortical topography. More broadly, these results illustrate the value of deriving the representational model space directly from neural data. Encoding models and representational similarity analyses have provided important insights into ventral temporal cortex organisation^9,16-18,50^, but they inherit the assumptions of their chosen feature spaces and may overlook dimensions that are neurally prominent yet semantically or behaviourally underspecified. Ultra-high-field imaging combined with dense stimulus sampling^24^ provided the statistical power and representational breadth needed to estimate a continuous geometry across the full object space, and the recovered structure was stable under balanced category resampling, highly consistent across individual participants, and replicated in two independent datasets acquired with different scanners, tasks and designs. Inverting the standard encoding-model logic in this way, by inferring the representational model from neural data and interpreting it post hoc, is likely to become increasingly productive as high-resolution imaging and large-scale stimulus databases become more widely available. The geometry also offers a well-specified target for computational accounts. Topographic neural network models, in which smooth representational maps, category-selective clusters and cortex-like gradients emerge from spatial locality and wiring constraints alone, without explicit category supervision^51^, can now be evaluated against a three-poled arrangement with named anchors and a lateral-to-medial cortical implementation, and no longer against category-selectivity benchmarks alone.

Several limitations qualify these conclusions. First, varimax rotation fixes the orientation of the axes within the leading subspace according to a simplicity criterion, so that the subspace, the triangular arrangement within it and its cortical mapping are invariant to this choice, whereas the specific labelling of Factor 1 and Factor 2 is not. Our claims accordingly rest on the geometry and not on any privileged status of the two axes. Second, exploratory factor analysis is a linear method ^52,53^, so any nonlinear structure in the representational manifold falls outside its scope, and nonlinear dimensionality reduction or more flexible latent-variable models^22^ would be a natural extension that could reveal additional geometric complexity without sacrificing interpretability. Third, several of the behavioural dimensions most strongly aligned with the axes describe appearance, and our region of interest extended into intermediate ventral visual areas; thus, cross-dataset agreement cannot by itself exclude a contribution of mid-level image statistics. We regard this as a bound on interpretation instead of a confound to be eliminated, given that the ventral temporal cortex is unlikely to separate cleanly into visual and conceptual codes^13,34^, although explicit comparison against image-computable feature models remains an important next step. Fourth, the orientation and semantic interpretation of the axes may depend on the breadth of the sampled stimulus space. Although the ecological diversity of the THINGS database was well suited to exposing this structure, our main analyses were restricted to the 960 concepts carrying top-down WordNet labels, and broader and denser sampling including dynamic stimuli, naturalistic scenes and multimodal object representations^43^ will be needed to test how far the geometry generalises. Finally, the present sample comprised twenty healthy young adults, a design that prioritises depth of sampling within individuals over breadth across the population, leaving open whether the geometry varies with age, culture or visual experience. Linking the higher-order factors, which carry reliable information about finer-grained distinctions, to behaviour^50^ is a further priority.

Notwithstanding these limitations, these results establish that the apparent complexity of object representations in the human ventral temporal cortex is organised by a small number of robust, interpretable axes recoverable directly from neural data. The resulting geometry, implemented as a smooth cortical gradient, accommodates both the localised selectivity seen in univariate studies and the broad decodability revealed by multivoxel approaches, and points towards a unified understanding of biological vision and the representational structures that emerge in computational models of visual processing.

## Data availability

The raw neuroimaging and behavioural data generated in this study cannot be made openly available because of restrictions imposed by the local institutional ethics approval and the informed consent obtained from participants. De-identified individual-level derivatives, including trial-wise fMRI response estimates, will be shared with qualified researchers under restricted access, contingent on approval of a data-use agreement and compliance with institutional ethics guidelines. Access requests should be submitted to the corresponding author and will be reviewed by the institutional data access committee. Requests will typically receive a response within four weeks, and approved data will be made available for non-commercial research for five years following approval. Group-level statistical maps will be made publicly available on the OSF repository (accession to be provided upon acceptance). The THINGS stimulus database is publicly available at https://things-initiative.org.

## Code availability

All original code generated for this study is publicly available as of the date of publication at https://github.com/cognizelab/neucognize. The experimental task paradigms were implemented using PsychoPy software (v2021.1.4). Preprocessing followed the Human Connectome Project (HCP) minimal preprocessing pipelines as implemented in QuNex (https://qunex.readthedocs.io), Singularity container. Cortical regions were denoted using the Multi-modal Parcellation Atlas (HCP_MMP V1.0). Cortical maps were visualised using Connectome Workbench (https://www.humanconnectome.org/software/connectome-workbench).

## Acknowledgements

We thank Peixin Yang and Yueting Su for assistance with data acquisition. We thank Kendrick Kay (University of Minnesota) for guidance on experimental design and for helpful discussions. We also thank the staff at the Zhangjiang International Brain Imaging Centre (ZIC) for support during data collection; Ying-Hua Chu (MR Research Collaboration Team, Siemens Healthineers Ltd.) for assistance with protocol setup; and Matthew F. Glasser (Washington University School of Medicine in St. Louis) and Essa Yacoub (University of Minnesota) for advice on optimising HCP-style data acquisition protocols.

## Author Contributions Statement

Y.W. and D.V. contributed to the conception and design of the work, and collected the data. The data were analysed by Y.W., X.L., K.Z., M.H., J.F. and D.V. The first draft was written by Y.W. and D.V. All authors contributed to revision of the manuscript.

## Methods and Materials

### Participants

All experimental procedures complied with the Declaration of Helsinki, Ethical Principles for Medical Research Involving Human Participants, and were approved by the local institutional review board (AF/SC-18/20210604). Twenty healthy young adults were recruited from the university population, according to predefined inclusion and exclusion criteria (mean age = 24.56 years, SD = 2.42, range = 19-29 years, female/male = 14/6). All participants were right-handed, native Mandarin speakers, had normal or corrected-to-normal vision, and reported no prior history of neurological or psychiatric illness and no contraindications to MRI. Following a detailed explanation of the study objectives and the multi-session dense sampling protocol, participants provided written informed consent and received monetary compensation (100 RMB/hour). Sample size was guided by prior studies employing similar dense sampling approaches^24,43^.

### Experimental procedure

Participants completed six visits at the Zhangjiang International Brain Imaging Centre (ZIC), Fudan University. During their first visit, we acquired 3T anatomical (T1w, T2w) and resting-state fMRI data. The remaining five visits were allocated for 7T task fMRI, conducted on weekdays (Monday-Friday), with scan times kept as consistent as possible across days. The median interval between consecutive sessions was 23.8 hours. During each visit, participants completed the main object recognition task and a functional localiser paradigm. Visual analogue scales assessing cognitive and affective states were collected immediately before and after each 7T session. Following the final 7T visit, participants also completed a set of post-scanning auxiliary behavioural assessments (not analysed here), including a standard neuropsychological battery, a multi-arrangement task, and an eye-tracking experiment.

#### Object recognition task

The main experimental paradigm, namely the object recognition task (ORT), was adapted from a continuous recognition paradigm used in a prior dense sampling experiment^24^. Stimuli were drawn from the THINGS image database^23^, comprising 22,256 high-quality, naturalistic object images embedded within contextual backgrounds, spanning 1,854 object concepts. The breadth of object concepts and multiple exemplars per concept included in this image set directly support analyses of object representations in the human brain that generalise across image-specific variation. For the present study, we focused on a subset of 960 concepts for which top-down WordNet labels were available. These labels were manually merged into 14 high-level semantic categories for category-level visualisation and analyses (animal, body part, clothing, decoration, weapon, food, plant, musical instrument, sports equipment, toy, tool, container, vehicle, and furniture; Table S1).

On each trial, a single object image was presented on a uniform grey background while participants maintained central fixation on a transparent red dot. The objective of the task was to indicate by button press whether the presented image was new or old (i.e., previously presented in the current or any prior session). Each trial lasted 4 s, in which the image was presented for 3 s, followed by a 1 s blank interval. Each run contained 75 trials: 64 image trials and 11 blank trials (three at the beginning, four at the end, and four interspersed pseudo-randomly among image trials). The interspersed blank trials were arranged so that the number of consecutive image trials ranged from 9 to 14. At the end of each run, participants received feedback indicating the number of responses recorded. Visual stimuli were presented onto a rear projection screen inside the bore with an MRI-compatible PRO-Pixx projector (VPixx Technologies Inc., Canada) and viewed via a mirror mounted on the head coil. Viewing distance was 202.5 cm (display resolution 1920 x 1080; refresh rate 120 Hz; image resolution 800 x 800; visual angle ∼6°). Responses were recorded using an MRI-compatible 4-button response box.

Across five 7T fMRI sessions, each participant completed 60 runs of the object recognition task paradigm (12 runs per session). We selected 22,256 images from THINGS to cover all 1,854 concepts (12 images for 1,846 concepts and 13 images for 8 concepts), specifically by taking image indices 1-12 for the 1,846 concepts and 1-13 for the remaining 8 concepts. The only a priori content-based exclusion criterion was the removal of images depicting threat objects e.g. firearms pointed directly at the viewer. No additional concept- or image-level exclusions were applied. These images were partitioned into 176 shared images that were common across participants, and 1,104 unique images per participant totalling 22,080 unique images. Each participant viewed 1,280 images in total (1,104 unique + 176 shared), and each image was presented three times to any given participant (3,840 image trials per participant).

Following prior work^24^, the ordering of the 3,840 image trials was pre-determined and identically applied to all participants. Specifically, the presentation of images was generated using a mixed distribution (60% von Mises with κ = 10; 40% uniform). Shared-image trials were assigned to the same slots for all participants, whereas the remaining unique-image slots were filled with participant-specific unique images, thereby maintaining a balanced mixture of novel and repeated images across runs and sessions.

#### Functional localiser task

After completion of all 12 object recognition task runs in each 7T fMRI session, participants performed a validated functional localiser task^54^ to independently identify tuned responses in the ventral temporal cortex. Grayscale images from ten categories were grouped into five stimulus domains: faces (adult, child), body parts (whole body, limb), places (corridor, house), objects (car, instrument), and characters (pseudoword, number), and displayed on scrambled backgrounds. Each 6 s block comprised 12 rapid serial visual presentations (500 ms per image) drawn from a single category. Each run included six repetitions per domain, interspersed with blank trials. Participants maintained central fixation and performed an oddball detection task by responding to trials containing scrambled images. The standard stimulus categories (body, word, adult face, car, house) were used in sessions 1, 3, and 5. Alternative categories (limb, number, child face, instrument, corridor) were used in sessions 2 and 4, adding to five runs in total across five sessions.

### MRI data acquisition

Data acquisition followed Human Connectome Project (HCP) guidelines^55^. Anatomical and resting-state scans were acquired on a 3T Siemens Magnetom Prisma (Siemens, Erlangen, Germany) with a 32-channel head coil using protocols adapted from HCP-Lifespan^56^. Task fMRI data were acquired on a 7T Siemens Magnetom Terra equipped with a 1-channel transmit/32-channel receive (1Tx/32Rx) head coil (Nova Medical, Wilmington, MA, USA) using harmonised sequences from the UK-7T network^57^. Compliant with HCP preprocessing pipelines^58^, this dual scanner protocol provided an optimal balance between whole-brain anatomical coverage with minimal distortions and high-resolution functional mapping of task-evoked brain activity.

At 3T, whole-brain anatomical imaging included a T1-weighted MPRAGE sequence (0.8 mm isotropic resolution; TR = 2500 ms; TE = 2.22 ms; TI = 1000 ms; flip angle = 8°; bandwidth = 220 Hz/pixel; in-plane acceleration factor [iPAT] = 2; acquisition time [TA] = 6 min 40 s) and a T2-weighted SPACE sequence (0.8 mm isotropic; TR = 3200 ms; TE = 563 ms; bandwidth = 745 Hz/pixel; iPAT = 2; TA = 5 min 57 s). Resting-state fMRI data (eyes open, fixating on a crosshair) were acquired using a multiband gradient-recalled echo echo-planar imaging (MB-GRE-EPI) sequence (2.0 mm isotropic; TR = 800 ms; TE = 37 ms; flip angle = 52°; bandwidth = 2290 Hz/pixel; multiband factor = 8; 488 volumes; TA = 6 min 40 s), with both anterior-posterior (AP) and posterior-anterior (PA) phase encoding directions to facilitate post-hoc distortion correction. Additional spin-echo and dual-echo gradient field maps were collected for geometric distortion correction.

Task-based fMRI at 7T employed a comparable multiband GRE-EPI sequence optimised for high spatial resolution (1.5 mm isotropic; TR = 1500 ms; TE = 25 ms; flip angle = 65°; multiband factor = 4; phase encoding = AP; TA = 8 min 40 s). Matching dual-echo gradient field maps were acquired before every four task runs to correct for susceptibility-induced distortions.

### MRI preprocessing

Neuroimaging data were preprocessed using HCP minimal preprocessing pipelines^58^ implemented in the Quantitative Neuroimaging Environment & Toolbox (QuNex)^59^. Structural images were corrected for gradient nonlinearity and aligned to the MNI space. Tissue segmentation and cortical surface reconstruction were performed via FreeSurfer^60^. Functional data were corrected for gradient distortions, head motion, and EPI susceptibility distortions, registered to the MNI152 template, and were projected to the standard 32k fsLR grayordinate space. The resulting dense timeseries comprised 91,282 grayordinates (59,412 cortical vertices and 31,870 subcortical voxels). No temporal filtering, detrending or additional smoothing, beyond the native 2 mm FWHM, was applied. In order to optimise the alignment with trial timings in the main object recognition task, preprocessed BOLD fMRI timeseries were upsampled to 1 s intervals using *tseriesinterp* function^61^.

### Trial-specific neural response estimation

Trial-specific neural response amplitudes for the object recognition task were estimated using GLMsingle^25^, which extends standard general linear model (GLM) analysis through three optimisation steps: (1) selecting an optimal hemodynamic response function from a library of 20 candidate HRFs, (2) deriving data-driven nuisance regressors via GLMdenoise^62^ (estimated across runs within each scan session), and (3) regularising single-trial beta estimates using fractional ridge regression. The analysis was performed in 32k fsLR surface space using the fully optimised betas_fithrf_GLM-denoise_RR output (incorporating all three GLMsingle components). Single-trial beta estimates were z-scored within each scan session to account for non-stationarities and standardise units across grayordinates, and were then averaged across scan repetitions to generate image-specific response profiles. For higher-level analyses, image-level betas were aggregated by THINGS object concept (pooling all available trials/images for that concept) and then averaged within 14 high-level semantic categories to derive category-level response patterns.

### Data quality assessment

Quality of the densely sampled task fMRI data was first assessed based on task performance. For that purpose, behaviour in the object recognition task was summarised using response rate and an adjusted hit-rate measure of recognition accuracy. Response rate was defined as the percentage of trials with a valid response (old or new), averaged within run and then across sessions. To assess recognition performance, trials were classified using standard signal-detection definitions: hits (old response to old item), misses (new response to old item), false alarms (old response to new item), and correct rejections (new response to new item). Hit rate was computed as hits/(hits + misses), and false-alarm rate as false alarms/(false alarms + correct rejections). Recognition accuracy was quantified based on the adjusted hit rate (hit rate − false-alarm rate).

For the assessment of session specific effects on behavioural performance, we fitted linear mixed-effects models with session as a fixed effect and participant as a random intercept. For response rate and adjusted hit rate, nested model comparisons were performed using likelihood-ratio tests between a full model (including session) and a reduced intercept-only model. To test whether recognition performance was above chance, we compared participants’ mean adjusted hit rate against zero using a one-sided one-sample t-test (alternative hypothesis: adjusted hit rate > 0), and we also report Cohen’s d. Unless otherwise stated, all other reported p-values were two-tailed with a significance threshold of 0.05.

In order to assess the quality of neuroimaging data, a comprehensive set of quantitative metrics were employed including: (1) temporal signal-to-noise ratio, (2) framewise displacement, (3) task-evoked BOLD response reliability, (4) noise ceiling estimation, and (5) cross-participant response consistency. Calculated as the mean signal divided by the temporal standard deviation for each run, session, and participant, temporal signal-to-noise ratio (tSNR) quantifies the capacity of fMRI acquisitions to detect neuronally-driven signal fluctuations. Framewise displacement, on the other hand, is a commonly used proxy measure of head motion, which is computed as the sum of the absolute derivatives of six rigid-body motion parameters (translations x, y, z; rotations α, β, γ). Following prior work^63^, rotational parameters were converted to displacement distances using the arc length on a sphere with radius 50 mm. No framewise displacement cut-off was applied, and no runs or sessions were excluded based on motion. Task-evoked signals were characterised using a basic ON-OFF GLM analysis. All stimulus trials were modelled as a single condition convolved with a canonical HRF. For each voxel, the variance-explained metric (ON-OFF R^2^) was defined as one minus the ratio of the residual sum of squares to the total sum of squares of the voxel-wise BOLD time series, thereby quantifying the proportion of temporal variance accounted for by the task model. Metrics from this model identified grayordinates carrying task-related signals and provided a quantitative measure of experiment-driven neural responses.

Participant-specific noise ceilings were computed following prior reports^24^ under four assumptions: (1) neural responses at each vertex/voxel depend solely on the presented image; (2) signal variability across images follows a Gaussian distribution; (3) additive noise is normally distributed with zero mean; and (4) observed responses represent the linear superposition of signal and noise components. Under these assumptions, the measured BOLD signal at each grayordinate follows a compound distribution resulting from the convolution of signal and noise distributions:

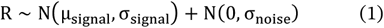

where R is the trial-wise beta response, µ_signal_ and σ_signal_ are the mean and standard deviation across different images, and σ_noise_ denotes standard deviation of the noise.

After z-score normalisation of trial-wise beta response estimates within session, image variance was calculated from repeated trials of the same image across all sessions. Standard deviation of the noise σ_noise_ was then estimated as follows:

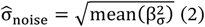

where 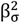is the image variance. Standard deviation of the signal was computed from 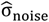 as follows:

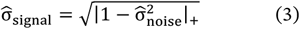

where | |_+_ is positive half-wave rectification. Next, the noise ceiling SNR ρ was computed as:

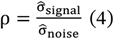

Finally, the noise ceiling across all sessions was computed as:

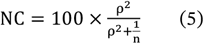

where n = 3 is the number of repeated trials (i.e., three presentations of the same image across the full experiment within participant).

Cross-participant consistency of neural responses to the 176 shared stimuli was quantified using the intraclass correlation coefficient (ICC). For each vertex, participants were treated as raters, shared images as targets, and response estimates as ratings. Specifically, we calculated the ICC(3,k) variant with k = 20, corresponding to a two-way mixed-effects, consistency, average-rating model. Cross-participant consistency of category-level representational geometry was assessed using representational similarity analysis (RSA)^48^ on the 176 images shared across all participants (see details below). Pairwise Pearson correlations were computed between all participant-specific representational dissimilarity matrices (RDM) across their 91 unique upper-triangular elements, averaged in Fisher-z space and back-transformed to Pearson’s r. Uncertainty in this mean was estimated using a participant-cluster bootstrap with 100,000 resamples, in which participants were sampled with replacement and repeated draws were represented using multiplicity weights, with the percentile distribution defining the 95% confidence interval. Within-participant RDM reliability was estimated from 1,000 balanced complementary allocations of the three presentations of each shared image, assigning one presentation to one half and two to the other half with singleton assignments balanced across categories and halves, and corrected to the full three-presentation estimator using the Spearman-Brown formula. Standard RSA noise ceilings were estimated from the participant-specific RDMs, with the lower bound obtained by correlating each participant’s RDM with the mean RDM of the remaining participants and the upper bound using the group-mean RDM including that participant.

### Neuroimaging data analysis

In order to characterise the neural representation of object concepts in the ventral temporal cortex, we combined exploratory factor analysis, multi-voxel pattern analysis and representational similarity analysis to provide a comprehensive view of how the brain organises objects representations.

#### Ventral-temporal-cortex definition

The ventral temporal cortex region of interest (ROI) combined participant-specific functional localiser responses, literature-based group masks and a multi-modal parcellation of the human brain (HCP-MMP). All ROI components were defined in the standard 32k fsLR surface space, giving vertex-wise correspondence across participants. Functional localiser data were analysed with a standard GLM. The first-level design included five boxcar regressors modelling each stimulus domain (faces, bodies, places, objects, characters), convolved with a canonical HRF, with pre-whitening to account for temporal autocorrelation^54,64^. Within participant, fixed-effects analyses combined runs, and group-level mixed-effects inference used non-parametric permutation testing (5,000 iterations) with PALM^65^. Domain-selective contrasts (each domain vs all others) were thresholded at the group level with FDR correction (q < 0.05). The union of significant category-selective vertices was then combined with literature-based masks of the ventral visual cortex^66^ and the multi-modal HCP-MMP atlas parcels from the secondary visual cortex to form the final region of interest within the ventral temporal cortex^67,68^. The final ventral temporal cortex mask comprised 12 parcels from the HCP-MMP corresponding to higher ventral visual areas: FFC, PH, PHA1, PHA2, PHA3, PIT, TE2p, V8, VMV1, VMV2, VMV3, and VVC.

#### Exploratory factor analysis

Exploratory factor analysis was used to identify latent dimensions underlying object representation in the ventral temporal cortex. This method constitutes a generative model of covariance, in which observed responses are expressed as linear combinations of a small number of latent factors plus residual (unique) variance^52^. This separation of shared and unique variance is advantageous when responses are noisy and when the goal is to recover dimensions that explain common structure across variables instead of simply maximising total variance.

In order to ensure that factor analysis was not biased by differences in mean activation or scaling, the matrix of trial-wise betas was z-score normalised for each vertex within session and then averaged across scan repetitions to generate image-specific responses. Data were preprocessed by excluding images shared across participants and averaging the remaining image-level responses within each object concept. No vertex in the ventral temporal cortex was excluded from analysis. For interpretability, only the subset of 960 object concepts carrying top-down WordNet labels was chosen for this analysis, resulting in a concept-by-vertex response matrix, denoted as Y. Suitability of the data matrix for factor analysis was evaluated using the Kaiser-Meyer-Olkin (KMO) measure and Bartlett’s test of sphericity. The KMO measure quantifies whether variables share sufficient common variance for factor analysis, and Bartlett’s test whether the correlation matrix differs from an identity matrix. Factor analysis was then performed using varimax rotation. The number of factors was selected by split-half cross-validation over 100 iterations with different random seeds, evaluating models with 10-100 factors (step size = 10). Average held-out log-likelihood peaked at 50 factors, and a 50-factor solution was adopted. In order to quantify the variance accounted for by the factors, an unrotated factor analysis model was also fitted and explained variance computed. The variance percentages reported in the main text derive from this unrotated solution, whereas all interpretation is based on the varimax-rotated factors. The two dimensions contributing most to the overall explained variance were chosen as the main representational space for further analysis.

### Functional and anatomical characterisation of the representational geometry

Following the identification of the two-dimensional representational geometry underlying object representation in the ventral temporal cortex, we next characterised the geometry of the two-dimensional latent space and the functional interpretation of the two factors.

#### Representational similarity analysis

Representational comparisons were performed on 14 x 14 category-level RDMs, constructed, compared and evaluated as described below.

#### Construction and comparison

Concept-level responses were averaged within the 14 high-level semantic categories. Vertex-space RDMs were computed as the Pearson correlation distance between category-level response patterns, and two-dimensional factor-space RDMs as the Euclidean distance between category-mean scores on Factors 1 and 2. Each RDM was summarised by its 91 unique upper-triangular elements, and any two RDMs were compared by the Pearson correlation across these elements.

#### Reliability

The reliability of an RDM was estimated by partitioning participants into two halves, constructing the RDM independently within each half, and correlating the two estimates across their 91 elements. Correlations were Fisher-z averaged, back-transformed to Pearson’s r, and corrected to the full-sample estimator using the Spearman-Brown formula:

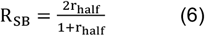

For the present dataset we used 200 balanced partitions of the 20 participants into groups of ten, with analyses restricted to concepts represented in both halves (957-960 concepts across partitions). When evaluating factor-space RDMs, a separate 50-factor varimax model was fitted within each participant half or participant, matched to the reference solution, and sign-aligned before the RDM was constructed.

#### Conditional fixed-reference ceilings

Where one RDM served as a fixed reference, namely the two-factor RDM in the full-sample comparison and the main-dataset RDM in all held-out and cross-dataset comparisons, the ceiling was defined as the square root of the Spearman-Brown-corrected reliability of the RDM being evaluated, and percent of ceiling as 100 x r_observed_ / r_ceiling_, calculated from unrounded values.

#### Uncertainty and significance

Confidence intervals for individual RDM correlations were obtained using a category-node bootstrap with 100,000 resamples, in which the 14 category nodes were sampled with replacement, only pairs involving different original categories were retained, and multiplicity weights were applied when computing the correlation. The 2.5th and 97.5th percentiles defined the 95% interval. For held-out and cross-dataset comparisons, significance was assessed using 100,000 category-label permutations, with the same permutation applied simultaneously to the rows and columns of the evaluated RDM so as to preserve symmetry and the dependence structure among pairwise distances, and one-sided p values calculated as p = (n_exceed_ + 1) / (n_perm_ + 1). These intervals quantify sensitivity to the sampled category nodes conditional on the fitted RDMs, and for external comparisons additionally on the two factor solutions and their matching.

#### Factor-space visualisation

Each concept was projected according to its scores on Factor 1 and Factor 2, and category-level distributions were obtained by grouping concepts within the 14 high-level semantic categories, with 95% confidence regions estimated for each category. This allowed us to assess whether categories formed discrete clusters or occupied overlapping positions within a continuous geometry.

#### Silhouette score

Cluster separation in the two-dimensional factor sub-space was quantified using the Silhouette score:

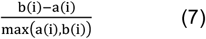

where a(i) denotes the mean Euclidean distance between concept i and all other concepts from the same category, and b(i) denotes the smallest mean

Euclidean distance between concept i and concepts from any other category. Values closer to 1 indicate well-separated clusters, values near 0 overlapping clusters at category boundaries, and negative values that concepts are on average closer to another category than to their own.

#### Multi-voxel pattern analysis

Linear support vector machine (SVM) classification was used to decode high-level object categories from vertex-wise response patterns or factor scores^49^. We utilised the LinearSVC implementation (based on the LIBLINEAR library) with regularisation parameter fixed at 1.0, which employs a coordinate descent solver to minimise the squared hinge loss. Classification performance was estimated using repeated split-half cross-validation, with data stratified by category, each half used in turn for training and testing, and accuracy averaged across the two folds. Significance was assessed against a label-shuffled baseline using Welch’s t-test.

For the main analysis of the two-dimensional factor space, classification focused on the three categories anchoring the triangular geometry (animals, foods and furniture). Balanced datasets were generated by sampling equal numbers of concepts from each of the three categories, and linear SVM classification was performed using their Factor 1 and Factor 2 scores. Estimates were averaged over 30 independently sampled balanced datasets, each with 10 random repeated split-half classifications.

For supplementary parcel-wise decoding, we performed one-vs-rest classification on vertex-wise response patterns within each ROI parcel, with balanced datasets sampling equal numbers of target and non-target concepts. Within each parcel, decoding accuracies were averaged across balanced resamples and repeated split-half classifications before visualisation.

### Semantic correlation analysis

With the aim of interpreting the semantic content of the latent dimensions, we correlated factor scores with 66 continuous semantic dimensions derived from large-scale behavioural similarity judgements in the THINGS database^42^. Pearson correlations were computed separately between each factor score and each semantic dimension across the 960 concepts. For each factor, semantic dimensions were ranked by correlation magnitude, and the strongest positive and negative associations were used to support interpretation of the latent axes. This analysis provided an independent behavioural reference for evaluating whether the neurally derived dimensions corresponded to interpretable object properties.

### Cross-validated prediction of factor scores

For assessing whether previously established object dimensions could account for the latent axes, we generated out-of-fold predictions of the first two varimax-rotated factor scores across the 960 concepts using nested cross-validated ridge regression. Three predictor sets were evaluated: binary animacy and THINGSplus real-world size, the 66-dimensional THINGS behavioural embedding, and their combination. Predictors were standardised within each training fold. Ten-fold outer cross-validation was used to estimate prediction performance, while the ridge penalty was selected within each training set using fivefold inner cross-validation over 17 logarithmically spaced values from 10^5;^ to 10^;^. Cross-validated R^2^ was calculated separately for Factor 1 and Factor 2.

### Participant-held-out prediction of neural RDMs

In order to test whether the factor-derived geometry explained neural representational structure beyond the a priori models, we constructed category-level RDMs from the two-dimensional factor space (D), binary animacy and standardised real-world size (A), and the 66 standardised THINGS behavioural dimensions (T). Ordinary least-squares models with intercepts compared A + T with D + A + T. Generalisation was evaluated across 200 balanced 10-versus-10 participant partitions in both train-test directions. For each direction, the factor model, factor-derived RDM, and regression coefficients were estimated using only the training participants and evaluated against the neural RDM from the held-out participants. Cross-validated R^2^ was not clipped at zero, and the primary statistic was:

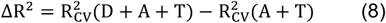

Percentile ranges across participant partitions were interpreted as summaries of split sensitivity.

### Generalisability and stability of latent representational geometry

Robustness was evaluated by repeating key analyses across data partitions, held-out stimuli and independent datasets.

Stability under concept resampling and across individuals. In order to assess sensitivity to category imbalance, we generated 100 category-balanced subsets by independently sampling 20 concepts from each of the 14 high-level semantic categories, giving 280 concepts per subset. When assessing consistency across individuals, we refitted the model separately for each of the 20 participants.

In both cases FA was refitted using the settings of the main analysis, and the resulting factor-loading matrices were aligned to the reference solution using the linear sum assignment algorithm to maximise the absolute Pearson correlation between loading vectors, followed by sign alignment. Similarity to the reference was then quantified in three ways: vertex-space category RDMs, two-dimensional factor-space category RDMs, and aligned factor-loading correlations for Factors 1 and 2. Correlations were Fisher-z averaged before back-transformation to Pearson’s r.

For the category-balanced subsets, confidence intervals for the expected mean under this fixed resampling procedure were estimated from 10,000 bootstrap resamples of the 100 subset-level estimates. These intervals are conditional on the observed participants, concept pool, preprocessing and analysis pipeline, and do not represent participant-population uncertainty.

#### Generalisation to held-out stimuli

The 176 images shared across all participants were excluded when fitting the reference FA model and were therefore treated as a held-out stimulus set. Vertex-space category RDMs were computed directly from their image-level responses after averaging within the 14 high-level categories. Factor-space category RDMs were computed by projecting the held-out vertex-wise response patterns through the fixed FA model estimated from the 960-concept dataset, without refitting, and taking Euclidean distances between the resulting category-mean scores on Factors 1 and 2. Both RDMs were compared with their counterparts from the full 960-concept dataset, treated as fixed references, using the correlation, reliability, ceiling, bootstrap and permutation procedures described under representational similarity analysis.

#### Cross-dataset generalisability

For comparing our results across independent datasets, we used neural response data from the THINGS-fMRI and CNeuroMod-THINGS datasets. The THINGS-fMRI dataset^43^ contains fMRI responses from three participants who completed 12 sessions while viewing 8,740 unique images spanning 720 THINGS object concepts. Images were presented in fast succession, participants maintained central fixation, and they performed an oddball-detection task in response to occasional synthetic images. The CNeuroMod-THINGS dataset^44^ contains data from four participants who completed 33-36 sessions of a continuous image-recognition task using THINGS stimuli from the same 720 concepts. For THINGS-fMRI, we used the released beta estimates in each participant’s native volume space, which were transformed to MNI space and then resampled to the 32k fsLR surface space. For CNeuroMod-THINGS, we started from the released preprocessed BOLD time series and estimated betas using GLMsingle, applying the same response-estimation approach used for the present dataset. Within each dataset, image-level responses were averaged within THINGS concepts and subsequently within the 14 high-level semantic categories.

To estimate the factor-space geometry, a 50-factor varimax-rotated FA model was fitted to each external dataset using the same model settings as in the main analysis. Factors were matched to the 50-factor reference solution estimated from the full 960-concept matrix Y using the linear sum assignment algorithm applied to the absolute Pearson correlations between factor-loading vectors. After factor matching and sign alignment, the components corresponding to reference Factors 1 and 2 were selected, and two-dimensional category RDMs were computed using Euclidean distances between category-mean factor scores. Cross-dataset similarity was quantified by correlating the 91 unique upper-triangular elements of the external-dataset RDM with those of the corresponding main-dataset RDM. Cross-dataset similarity was then quantified as described under representational similarity analysis. Factor-level correspondence was additionally quantified using absolute Pearson correlations between matched factor-loading vectors.

#### Latent-space response gradients

In order to characterise how neural response patterns vary continuously across the latent dimensions, we sampled coordinates along Factor 1 and Factor 2 separately in the two-dimensional factor space, while holding the non-target factor constant. Along Factor 1, the sampled coordinates were (-3,0), (-2,0), (-1,0), (0,0), and (1,0); along Factor 2, the sampled coordinates were (0,1), (0,0), (0,-1), (0,-2), and (0,-3). For each sampled coordinate, we computed its Euclidean distance to all 960 object concepts in the same two-dimensional space, identified the seven nearest concepts, and averaged their concept-level vertex-wise response profiles to estimate the local mean cortical response pattern. The resulting averaged response maps were then projected back onto the ventral cortical surface within the ventral temporal cortex.

#### Category-wise neural response gradients

For assessing category-wise response gradients, we computed mean neural responses for each of the 14 high-level categories within each ventral temporal cortex ROI parcel. Categories were ordered by their mean scores on Factor 1 or Factor 2. Parcels were ordered using a normalised FFA-PPA overlap index derived from independent probabilistic FFA and PPA maps^24^. These maps were obtained from the NSD fsaverage ROI probability maps and brought into the standard 32k fsLR surface space by first converting them to surface metric files and then resampling them with Connectome Workbench using area-preserving barycentric interpolation (ADAP_BARY_AREA) and the HCP fsaverage-to-fsLR spherical and midthickness templates. Within the ventral temporal cortex, vertices with positive overlap with the FFA or PPA localiser were identified, and for each parcel we computed the proportion of vertices overlapping with the FFA localiser minus the proportion overlapping with the PPA localiser. Positive values indicated relatively greater FFA overlap, whereas negative values indicated relatively greater PPA overlap. The resulting category-by-region response matrices were visualised as heatmaps to assess whether neural responses transition smoothly across cortical territory as a function of latent position.

