## Supplementary Information for "Two dominant axes structure high-dimensional object representations in the human ventral temporal cortex"

### Table of Contents

Supplementary Figures ........ 2
Supplementary Tables ........ 7

### Supplementary Figures

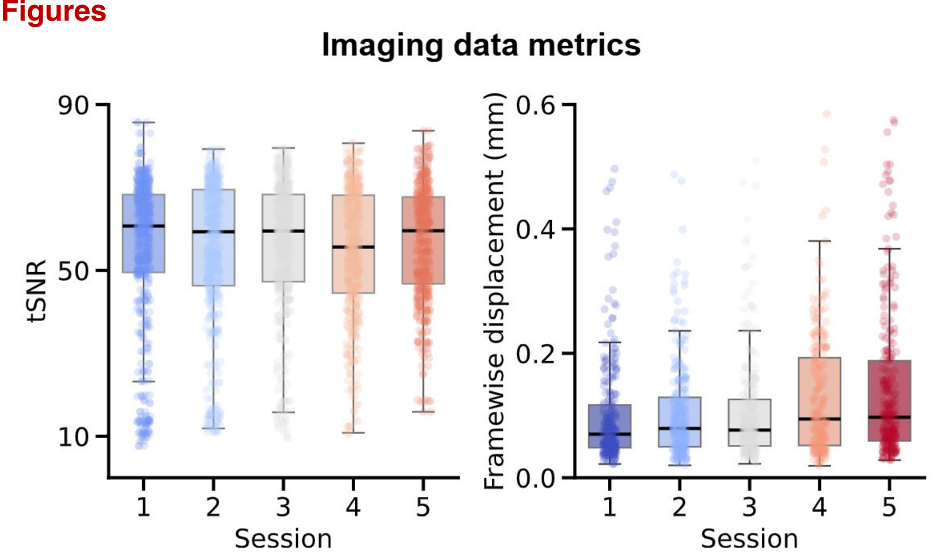


**Fig. S1 I Metrics of 7T fMRI data quality across scanning sessions.** The left panel shows temporal Signal-to-Noise Ratio (tSNR) and the right panel shows framewise displacement (mm). Each dot represents a single run from a participant's session, and boxplots indicate the distribution of data quality per session.

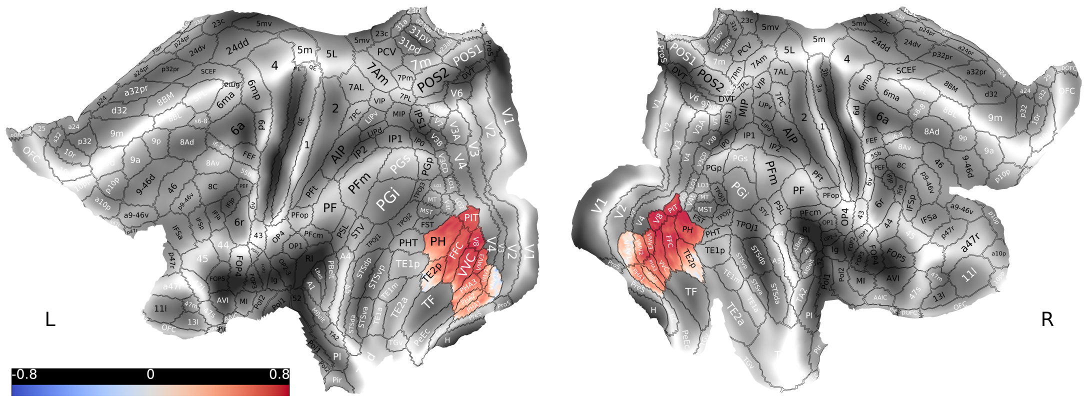


**Fig. S2 I Intraclass correlation coefficient (ICC) of shared images.** This figure displays the representational stability of 176 shared images across the ventral temporal cortex (VTC). Vertex-wise ICC values were computed using the shared stimuli as targets and participants as raters, providing a spatially resolved measure of cross-participant consistency. The resulting map highlights the degree to which shared-object responses were reliable across individuals in the densely sampled dataset.

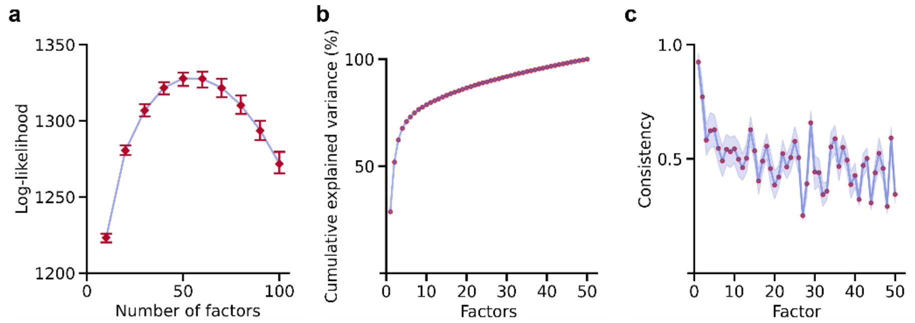


**Fig. S3 I Factor analysis of concept responses. a.** Determination of the optimal number of factors. We determined the optimal number of factors by running factor analysis on different number of factors on one half of the 960 concepts and computing the average log-likelihood of the other half. The optimal number of factors was chosen as the maximum log-likelihood. **b.** Cumulative explained variance by factors on the 960 concepts. The first few factors account for a large proportion of shared variance, with the remainder distributed across many further dimensions. **c.** Consistency of the factors on the 960 concepts. The plot shows how consistently each factor is recovered across random resampling iterations, demonstrating that the leading factors are highly reproducible.

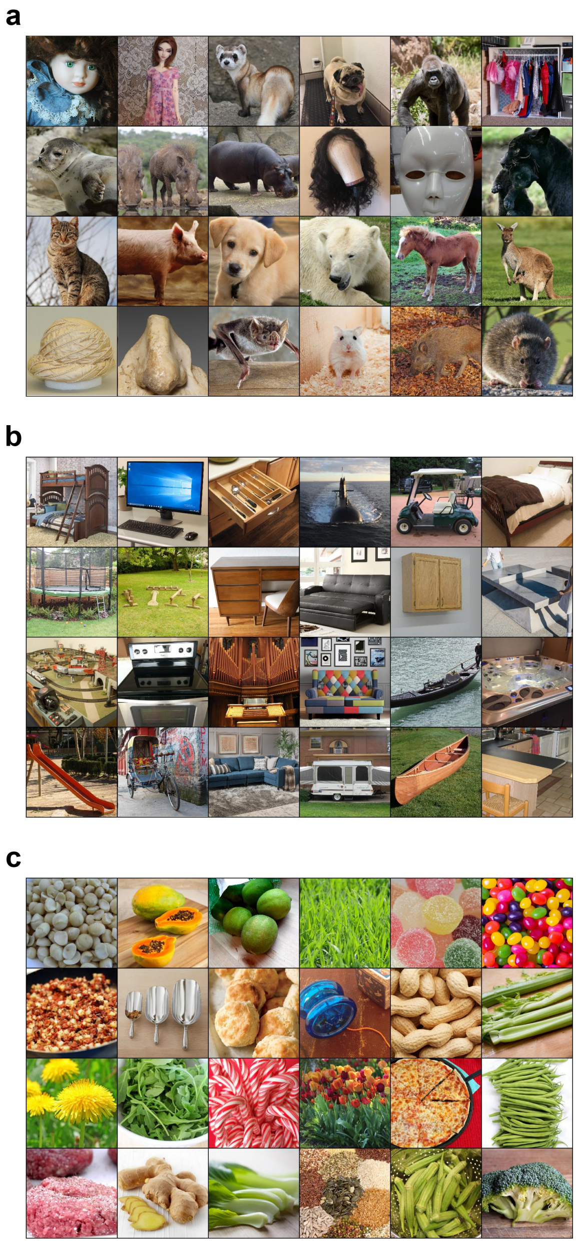


**Fig. S4 | Representative images of the concepts in the triangular poles. a.** Representative images of the concepts in the top-left of Fig. 3a inset. **b.** Representative images of the concepts in the bottom-right of Fig. 3a inset. **c.** Representative images of the concepts in the top-right of Fig. 3a inset. These examples illustrate the kinds of objects that anchor the main geometric poles of the representational manifold and help visualise how the latent dimensions map onto concrete object concepts. For privacy protection, several representative photographs containing real people or human body parts were replaced while retaining their original concept labels: the image for the face concept was replaced with a doll-face exemplar from THINGS; the image for the turban concept, which originally depicted a person wearing a turban, was replaced with a standalone turban; and the image for the nose concept was replaced with a non-identifying nose exemplar. The latter two replacement images were obtained from Wikimedia Commons under CC0 licensing. These substitutions are limited to figure visualization only.

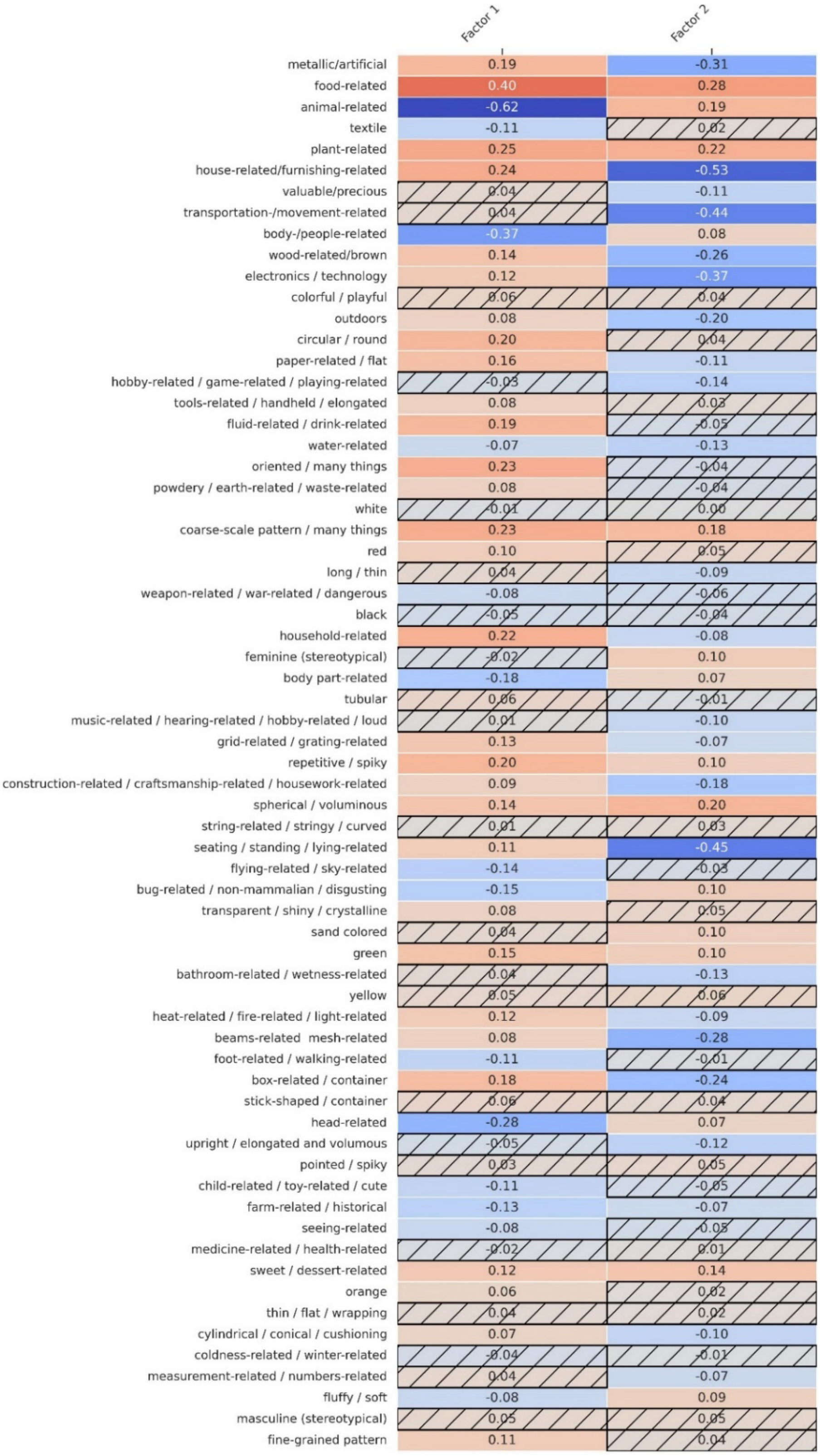


**Fig. S5 | Correlation matrix between 66 THINGS semantic dimensions and Factors 1/2.** The heatmap shows Pearson correlation coefficients: red (positive), blue (negative). Hatched cells mark insignificant correlations ($p > 0.05$).

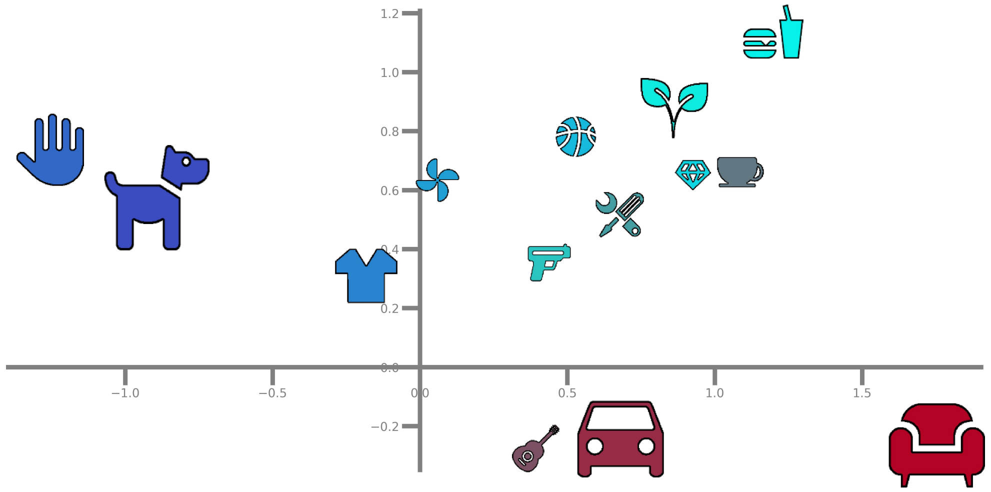


**Fig. S6 I Two-dimensional category scores of shared images.** This figure shows the projection of the 176 shared images into the learned two-dimensional factor space. The main latent geometry is preserved when analysis is restricted to the shared stimulus subset. The figure therefore provides a direct test of whether the recovered representational geometry generalises to stimuli not used in the primary concept-level analysis.

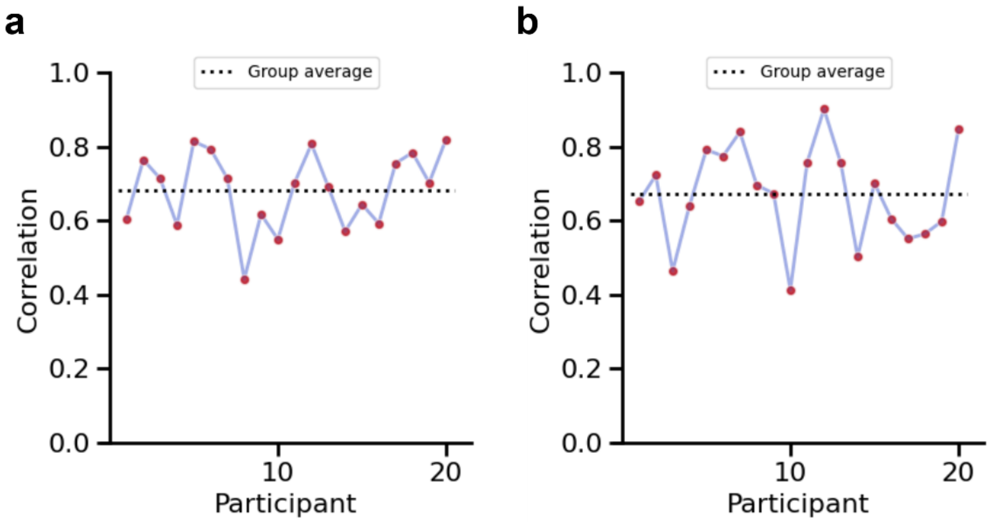


**Fig. S7 I Factor analysis of individuals. a.** Vertex-level categorical RDM correlations between the 960 object concepts and individuals. Black dash is mean across participants. **b.** Categorical RDM correlations between the 960 object concepts and individuals in two-dimensional subspace. Black dash is mean across participants. Together, these panels show that the latent geometry and its corresponding representational structure are recovered consistently at the level of individual participants.

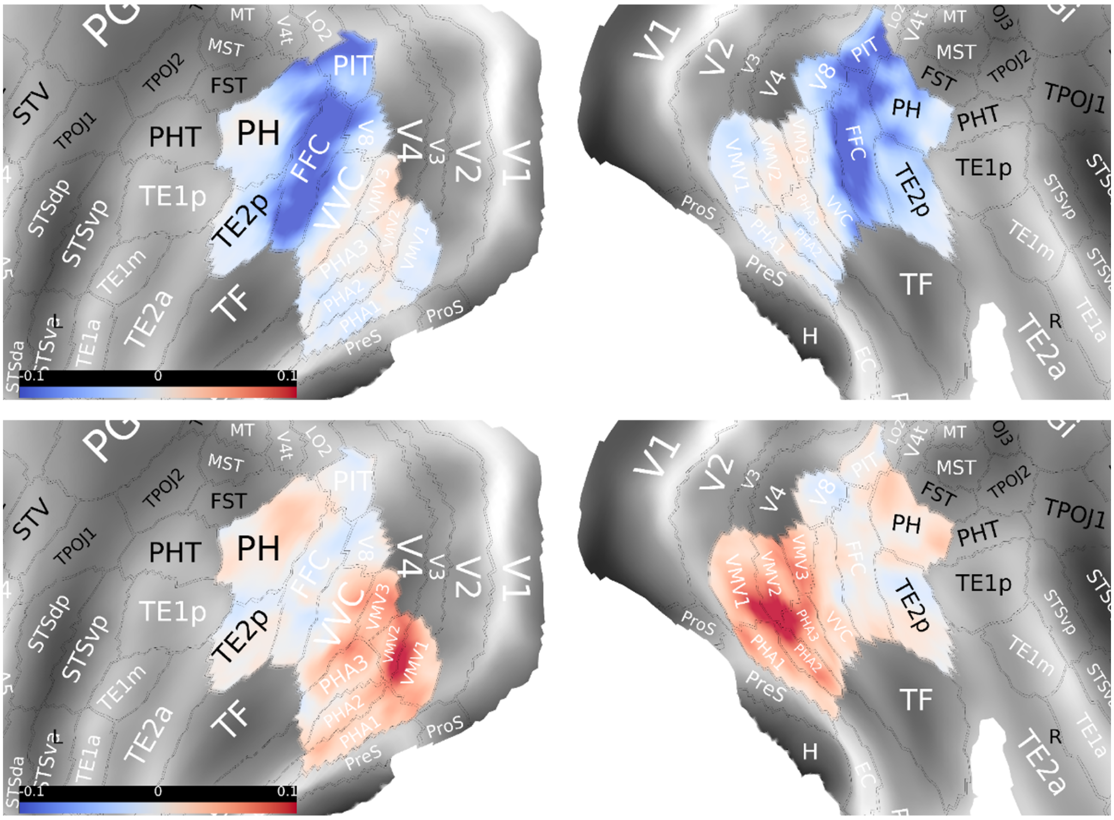


**Fig. S8 | Anatomical distribution of the first two factors of THINGS-fMRI.** This figure maps the factor loadings obtained from the independent THINGS-fMRI dataset back onto the cortical surface. The resulting spatial distribution shows whether the latent dimensions discovered in our dataset are also expressed as a similar cortical organisation in an external dataset acquired under different experimental conditions.

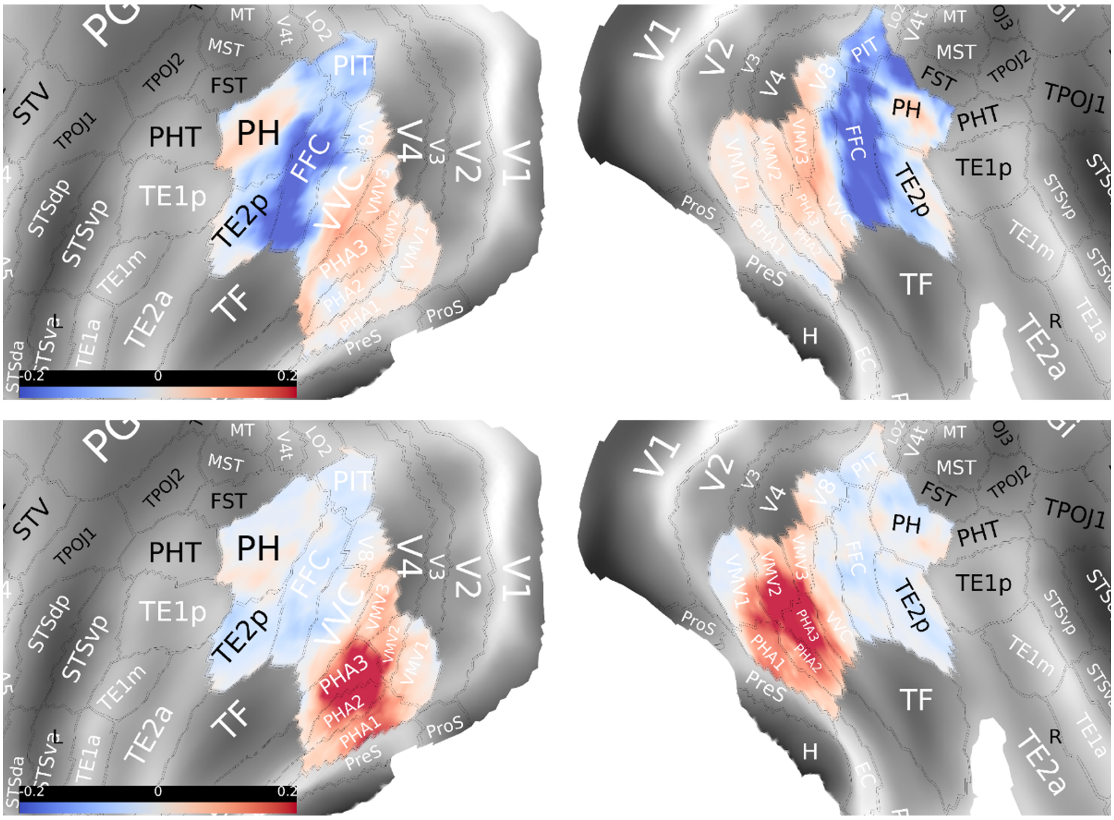


**Fig. S9 | Anatomical distribution of the first two factors of CNeuroMod-THINGS.** This figure shows the cortical projection of the first two latent factors in the independent CNeuroMod-THINGS dataset. The panel provides an additional cross-dataset test of whether similar factor loadings are recovered despite differences in scanner characteristics, participants, and task design.

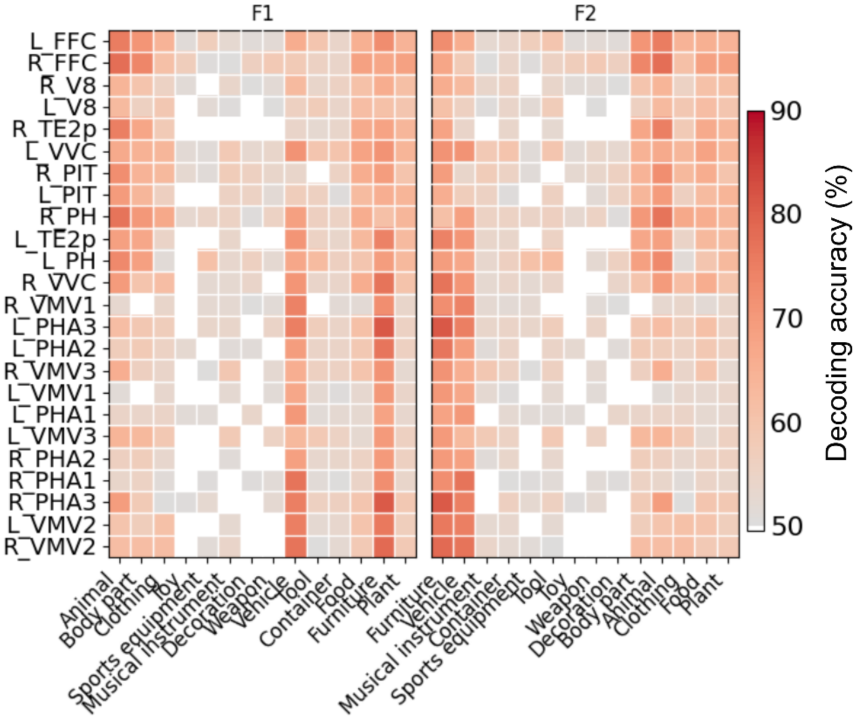


**Fig. S10 | ROI-wise category decoding accuracy across ventral temporal cortex parcels.** Heatmaps show mean binary one-vs-rest decoding accuracy for each of the 14 high-level object categories within each VTC ROI parcel. Rows denote parcels ordered by the normalised FFA–PPA overlap index used in Fig. 5c, progressing from parcels with stronger FFA overlap to parcels with stronger PPA overlap. Columns show the same categories ordered by their mean scores on Factor 1 (left) or Factor 2 (right). Values were computed from parcel-wise linear-SVM decoding and averaged across balanced resamples and repeated train-test splits. White colour indicates decoding accuracies that were not significantly above chance.

### Supplementary Tables

**Table S1 | Concept distribution across high-level semantic category and top-down WordNet labels.** This table summarises the number of object concepts included in each merged high-level semantic category, together with the corresponding top-down WordNet subcategories and their concept counts. It provides an overview of how the 960 THINGS concepts were distributed across the semantic categories used in the main analyses.

| No. | Merged high-level semantic category | Total concepts | Top-down WordNet category | Concept counts |
|---|---|---|---|---|
| 1 | animal | 174 | animal | 131 |
| | | | animal, bird | 27 |
| | | | animal, insect | 16 |
| 2 | plant | 110 | plant | 37 |
| | | | fruit | 49 |
| | | | vegetable | 24 |
| 3 | food | 151 | food | 121 |
| | | | food, dessert | 11 |
| | | | food, beverage | 16 |
| | | | food, vegetable | 3 |

| No. | Merged high-level semantic category | Total concepts | Top-down WordNet category | Concept counts |
|---|---|---|---|---|
| 4 | container | 100 | container | 96 |
| | | | container, kitchen utensil | 4 |
| 5 | vehicle | 65 | vehicle | 26 |
| | | | container, vehicle | 38 |
| | | | container, sports equipment, vehicle | 1 |
| 6 | tool | 78 | tool | 50 |
| | | | kitchen utensil | 13 |
| | | | kitchen appliance | 10 |
| | | | electronic device | 5 |
| 7 | weapon | 20 | weapon | 18 |
| | | | sports equipment, weapon | 1 |
| | | | vehicle, weapon | 1 |
| 8 | body part | 31 | body part | 31 |
| 9 | decoration | 30 | decoration | 30 |
| 10 | clothing | 93 | clothing | 93 |
| 11 | musical instrument | 32 | musical instrument | 32 |
| 12 | sports equipment | 20 | sports equipment | 20 |
| 13 | toy | 24 | toy | 24 |
| 14 | furniture | 32 | furniture | 32 |
